# Patient-derived Glioblastoma Stem cells transfer mitochondria through Tunneling Nanotubes in Tumor Organoids

**DOI:** 10.1101/2020.11.21.392597

**Authors:** Giulia Pinto, Inés Saenz-de-Santa-Maria, Patricia Chastagner, Emeline Perthame, Caroline Delmas, Christine Toulas, Elizabeth Moyal-Jonathan-Cohen, Christel Brou, Chiara Zurzolo

## Abstract

Glioblastoma (GBM) is the most aggressive brain cancer and its relapse after surgery, chemo and radiotherapy appears to be led by GBM stem cells (GSLCs). Also, tumor networking and intercellular communication play a major role in driving GBM therapy-resistance. Tunneling Nanotubes (TNTs), thin membranous open-ended channels connecting distant cells, have been observed in several types of cancer, where they emerge to drive a more malignant phenotype. Here, we investigated whether GBM cells are capable to intercommunicate by TNTs. Two GBM stem-like cells (GSLCs) were obtained from the external and infiltrative zone of one GBM from one patient. We show, for the first time, that both GSLCs, grown in classical 2D culture and in 3D-tumor organoids, formed functional TNTs which allowed mitochondria transfer. In the organoid model, recapitulative of several tumor’s features, we observed the formation of a network between cells constituted of both Tumor Microtubes (TMs), previously observed *in vivo*, and TNTs. In addition, the two GSLCs exhibited different responses to irradiation in terms of TNT induction and mitochondria transfer, although the correlation with the disease progression and therapy-resistance needs to be further addressed. Thus, TNT-based communication is active in different GSLCs derived from the external tumoral areas associated to GBM relapse, and we propose that they participate together with TMs in tumor networking.

## Introduction

Glioblastoma (GBM) is the most common and aggressive brain cancer which nowadays lacks understanding and resolutive therapeutic strategies. After surgery, patients undergo a mixture of chemo and radiotherapy[1], aiming to kill the remaining cancer cells at the edges of the resected region. Although these treatments have been proven to be effective in extending patients survival[2], lethal relapse from these peripheral regions occurs in 100% the cases. The elevated intra-tumoral heterogeneity seems to be at the origin of the relapse and particularly due to the presence of GBM stem cells (GSCs) that have been found to be the most resistant to treatments[3–6]. Moreover, it has been shown that post-surgical treatments can induce cellular plasticity and trans-differentiation resulting in more aggressive phenotypes[7]. How this occurs is still not clear, however it appears that intercellular communication in the tumoral context has a major role in the plasticity, survival and progression of many different types of cancer[8,9]. In particular, in the case of GBM, Winkler and colleagues have shown that patient-derived GSCs, xenografted into murine brains, are able to grow tumors where cells interconnect through membranous extensions and form a unique communicating network[10]. These protrusions called Tumor Microtubes (TMs) range in the microscale for their diameter (>1μm) and could extend for over 500 μm in length, creating a complex tumor cell network. TMs allow the propagation of ion fluxes through GAP-junctional proteins, such as Connexin43 (Cx43), providing a fast, neurite-like, communication between cancer cells. These extensions could also drive the repopulation of surgically-injured areas[11]. TM-connected cells resulted to be resistant to chemo and radiotherapy, and the resistance was lost following the inhibition of TM-inducers such as Cx43, GAP43 and TTYH[10–12].

Another mechanism of intercellular communication that has been recently proposed to facilitate tumor progression is represented by Tunneling Nanotubes (TNTs)[13,14]. TNTs are thin cellular extensions connecting distant cells observed in a wide variety of cellular and murine models as well as in *ex vivo* resections from human tumoral tissue[13]. They are membranous structures supported by an actin-based cytoskeleton and, differently from other cellular protrusions, including TMs (assumed to provide communication through GAP-junction), are open at both extremities, thus allowing cytoplasmic continuity between connected cells[15,16]. TNTs allow the transfer of various-sized cargos, such as small molecules (e.g. Ca^2+^ ions), macromolecules (e.g. proteins, nucleic acids) and even organelles (vesicles, mitochondria, lysosomes, autophagosomes, etc.)[17]. They appear to play a critical role in several physio-pathological contexts, as in the spreading of protein aggregates in various neurodegenerative diseases[18–22] or in the transmission of bacteria[23] and viruses[24,25] and, possibly, during development[26]. Functional TNTs have been shown in a variety of cancers using *in vitro* and *ex vivo* models [13] where they could be exploited as route for the exchange of material between cancer cells or with the tumoral microenvironment. As consequence of this transfer, cells can acquire new abilities as enhanced metabolic plasticity, migratory phenotype, angiogenic ability and therapy-resistance. In particular, the transfer of mitochondria has been related to all the previously mentioned features since they can provide energy and metabolic support to the cancer cells in displaying their aggressive features as observed in various cancers[14,27].

Few studies have reported TNT-like communication in GBM cells lines[28–30], suggesting that their presence and functionality could be induced/affected by the treatments, contributing to the tumoral progression and treatment-resistance [31,32]. However, no data on the role of TNTs are available in the context of a whole GBM tumor or in primary GSCs. This is likely due to the fragility of these connections and to the low-resolution images that can be obtained in the *in vivo* studies [10]. Whether in GBM intercellular communication is orchestrated exclusively by TMs or whether TNTs are also present and functional is still not known. Here, we investigate for the first time if TNTs can be formed between patient-derived GSCs and be exploited for exchange cargos using a quantitative approach. We used GSCs derived from the infiltrative region of the tumor, responsible for GBM relapse, thus representing a relevant model for the progression of the disease. In these cells we addressed TNT presence and functionality in both classical adherent cell culture as well as in 3D tumor organoids as well as the effect of radiotherapy on the TNT-mediated communication.

## Material and Methods

### Cell culture

GBM samples were processed as described by Avril et al. 2011[33]. GSLCs were cultured in suspension in DMEM-F12 (Sigma) supplemented with B27 (50x Gibco), N2 (100x Gibco) and 20 ng/ml of FGF-2 and EGF (Peprotech) at 37°C in 5% CO_2_ humidified incubators. Fresh medium was added to the cell culture every 2-3 days. All GSLCs were used for the experiments in this medium at less than 25 passages. Absence of alteration upon culture passages on the stemness phenotype was monitored by RT-qPCR. Absence of mycoplasma contamination was verified with MycoAlert™ Mycoplasma Detection Kit (Lonza). All methods were carried out in accordance with the approved guidelines of our institution.

### Lentivirus preparation and transduction

Lentiviral particles were produced in human 293T cultured in in Dulbecco’s Modified Eagle’s Medium (ThermoFisher), supplemented with 10% Fetal Bovine Serum (EuroBio) and 1% Pen/Strep (100x Gibco) at 37°C in 5% CO_2_ humidified incubators. Cells were plated at a 50-70% confluency the day before the transfection. Plasmids coding for lentiviral components, pCMVR8,74 (Gag-Pol-Hiv1) and pMDG2 (VSV-G) vectors, and plasmid of interest at a ratio of 4:1:4, respectively were transfected using FuGENE HD Transfection reagent according to manufacturer’s protocol. MitoGFP (pLV-CMV-mito-GFP) and mCherry (pLV-CMV-mCherry) plasmids encode respectively for a fragment of the subunit VIII of human cytochrome C oxidase fused with GFP, and for cytosolic mCherry under the Cytomegalovirus (CMV) promoter. Viral particles were concentrated using LentiX-Concentrator (TakaraBio) after 48 hours, and GSLCs were infected and tested for the expression of the fluorescent marker by flow cytometry at different time points to monitor expression stability.

### Tumor organoids preparation and culture

Tumor organoids were prepared according to the protocol published in Hubert et al. 2016[34] and cultured in Neurobasal medium (ThermoFisher) supplemented with B27 (50x Gibco), N2 (100x Gibco), 1% Pen/Strep (100x Gibco), 2 mM L-Glutamine (100x Gibco), 20 ng/ml of FGF, 20 ng/ml EGF (Peprotech) and 1.6 mL of GelTrex (ThermoFischer) at 37°C in 5% CO_2_ humidified incubators up to 23 days. Part of the cultured medium was removed and replaced with fresh one every 2-3 days.

### TNT counting

TNTs were identified according to the protocol of Abounit et al., 2015[35]. Cell were plated at the ideal cell density for the observation of TNTs (40000 cells/cm^2^ for both GSLCs). The adhesion surface was previously coated with laminin 10 μg/mL (Sigma) for at least 2 hours. GSLCs were fixed after 6 hours, to avoid excessive cell flattening on the coated surface. 15 minutes fixation in solution 1 (2% PFA, 0.05% glutaraldehyde and 0.2 M HEPES in PBS) followed other 15 minutes in solution 2 (4% PFA and 0.2 M HEPES in PBS) were performed at 37°C in order to preserve TNTs integrity[35]. Plasma membrane was labelled with fluorescent Wheat Germ Agglutinin (1:500 in PBS, Life Technologie) for 20 min at RT. Nuclei ware stained with DAPI (1:5000 Sigma-Aldrich) before mounting with Mowiol.

Tile confocal images were acquired with a Zeiss LSM 700 controlled by ZEN software. Optimal image stack was applied. The whole volume of the cells was acquired. All the images were processed using ICY software in order to manually count the number of TNT-connected cells. Cells connected through thin, continuous, phalloidin-positive connections were counted as TNT-connected cells.

### Immunofluorescence

Cell were seeded on glass coverslips at the TNT-density previously mentioned. Coverslips were coated with 10 μg/mL laminin (Sigma). Cells were fixed with 4% PFA for 20 minutes at RT. Quenching and permeabilization steps were performed using 50 nM NH4Cl solution and 0.1-0.2% Triton-X100, respectively. Primary antibodies were incubated in 10% FCS-containing PBS solution for 1 hour. Anti-αTubulin (1:1000 Sigma-Aldrich T9026), anti-TOM20 (1:500 Santa Cruz Biotechnology sc-11415) and anti-GAP43 (1:500 Cell signalling 8945S) were used. Cells were incubated for 45 minutes with secondary antibody anti-mouse and anti-rabbit Invitrogen Alexa 488, 564 or 647 antibodies (1:1000) or Rhodamine-conjugated Phalloidin (1:200, R415 Invitrogen). DAPI (1:5000 Sigma-Aldrich) in PBS solution was applied for 5 minutes before washes and mounting with Mowiol.

Organoids were fixed with a solution of 4% PFA for 1 hour at 37°C, washed with PBS-0.5% Tween and incubated in a solution of PBS + 10% FBS + 0.3% BSA (Sigma-Aldrich) + 0.3% Triton-X100 0.3% containing primary antibody (mentioned above) overnight at 4°C. Organoids were incubated in the same solution with the corresponding secondary antibody or Rhodamine-conjugated Phalloidin overnight at 4°C, incubated with DAPI (1:1000 Sigma-Aldrich) over 6h and finally mounted with a solution of 70% Glycerol.

Immunofluorescence stainings were analysed on a Zeiss LSM 700 inverted confocal microscope (Carl Zeiss, Germany), with a Pln-Apo 10X/0.45 to image the entire organoid, 40X: EC Pln-Neo 40X/1.3 (NA = 1.3, working distance = 0.21mm) or Pln-Apo 63X/1.4 (NA = 1.4, working distance = 0.19mm) oil lens objective and a camera (AxioCam MRm; Carl Zeiss).

### Time-lapse Microscopy

Time-lapse microscopy imaging in 2D- and 3D-conditions was performed on an inverted Spinning Disk microscope (Elipse Ti microscope system, Nikon Instruments, Melville, NY, USA) using 60 × 1.4NA CSU oil immersion objective lens using Bright field and Laser illumination 488. Pairs of images were captured in immediate succession with one of two cooled CCD cameras, which enabled time intervals between 20 and 30 s per z-stack. For live cell imaging, the 37 °C temperature was controlled with an Air Stream Stage Incubator, which also controlled humidity. Cells placed in Ibidi μ-Dish 35 mm and incubated with 5% CO_2_ during image acquisition. Image processing and movies were realized using MetaMorph, FIJI and Imaris software. Time-lapse movies of mitochondria trafficking were created using ImageJ/Fiji software.

### Quantification of mitochondria transfer by flow cytometry

Transfer assays were performed accordingly to the protocol of Abounit et al., 2015[35]. Stable GSLCs population expressing respectively MitoGFP were used as donor cells and mCherry as acceptor cells and mixed in a 1:1 ratio. For the 2D co-culture, cells were plated at the density previously mentioned (see TNT counting). Cells were detached after 2 or 5 days of co-culture with StemPro Accutase (Thermofisher), experimental duplicates were performed for each timepoint and each condition. To monitor the transfer by secretion in 2D co-culture, donor and acceptor cells were co-cultured separated by a 1 μm filter. For tumor organoids, donor and acceptor cells were mixed 1:1 during the organoid preparation. At each timepoint, duplicates of a pool of 3 organoids were disaggregated using mechanical and chemical (StemPro Accutase, Thermofisher) dissociation. To monitor the transfer by secretion, organoids prepared of only acceptor or donor cells were cultured in the same culture medium separated by a 1 μm filter. For FACS analysis, cells were passed through a cell strainer to separate cell aggregates and fixed in 2% PFA. Flow cytometry data were acquired with a BD Symphony A5 flow cytometer. GFP and mCherry fluorescence were analysed at 488 nm and 561 nm excitation wavelength, respectively. 10,000 events were acquired for each condition and data were analysed using FlowJo analysis software.

### The extracellular flux cell mitochondrial stress analysis (Seahorse assay)

An extracellular flux analyser (Seahorse XF96, Agilent, USA) was applied to analyse the mitochondrial function. The XF96 possesses a specialized microplate that allows for the measurement of the oxygen consumption rate (OCR) in real-time [36]. To test mitochondrial respiration, an XF Cell Mito Stress Test kit (Seahorse Bioscience; Agilent Technologies, Inc.) was used according to the manufacturer’s protocol. The day previous to assay, the cells were seeded at a density of 20,000 cells/well in a laminin-precoated seahorse plate and incubated overnight at 37°C and 5% CO_2_. The sensor cartridge was hydrated in pure water at 37°C in a non-CO_2_ incubator overnight. On the day of the assay, the sensor cartridge was incubated in XF Calibrant 1h at 37°C in a non-CO_2_ incubator prior to the assay. The culture medium was refreshed 1 h prior to the assays using unbuffered DMEM (pH 7.4) supplemented with 1 mM pyruvate, 2mM glutamine, and 10 mM glucose (Seahorse Bioscience; Agilent Technologies, Inc.). Briefly, 1 μM oligomycin, 1μM carbonyl cyanide-4-(trifluoromethoxy) phenylhydrazone (FCCP), and 0.5 μM rotenone/antimycin A were subsequently added to the microplates. This enabled determination of the basal level of oxygen consumption, ATP-linked oxygen consumption, non-ATP-linked oxygen consumption, the maximal respiration capacity, and the non-mitochondrial oxygen consumption. A total of three basal OCR measurements were recorded prior to the injection of oligomycin. The decreased level of OCR represented oligomycin-sensitive OCR due to its inhibition of ATP synthase (complex V). FCCP, an uncoupling protein, was then injected and the FCCP-stimulated OCR was used to calculate spare respiratory capacity, which was defined as the difference between maximal respiration and basal respiration. The third injection was a mixture of rotenone (a complex I inhibitor) and antimycin A (a complex III inhibitor). This combination inhibited mitochondrial respiration completely, and thus no oxygen was further consumed by cytochrome c oxidase. The remaining OCR measurement obtained following this treatment was primarily non-mitochondrial and may have been due to cytosolic oxidase enzymes.

### Irradiation

Irradiation was performed with X-Ray machine (Xstrahl LTD). 2 Gy irradiation was performed exposing the cells to X-rays for 1 minute and 25 seconds (250 kV, 12 mA).

### RT-qPCR

Total RNA extraction was performed using the RNeasy Mini Kit purchased from Qiagen. Reverse transcription was done using the Biorad iScript gDNA Clear cDNA Synthesis Kit. Oligonucleotides were designed using Prime PCR Look Up Tool (Bio-Rad), purchased from Eurofins Genomics, and sequences are presented in Supplementary Table 1. Quantitative PCR was then performed using the Bio-Rad iTaq™ universal SYBR^®^ Green supermix and analysed using a CFX96TM real-time PCR detection system under the CFX Manager software (Bio-Rad). Gene expression was normalized to hypoxanthine-guanine phosphoribosyltransferase (HPRT), each point of each independent experiment was performed in triplicates. Oligonucleotides used in qPCR are presented in Supplementary Table 1.

### Statistical analysis

The statistical tests for percentage of connected cells and percentage of transfer were computed using either a logistic regression model computed using the ‘glm’ function of R software (https://www.R-project.org/.) or a mixed effect logistic regression model using the lmer[37] and lmerTest[38] R packages. For cell connection in 2D, a mixed effect logistic regression model was estimated, adjusted on the effect of cell type, timepoint and condition. This model was also adjusted on the second and third order interactions among these 3 covariates. A random effect corresponding to replication of the experiment was also added to the model in order to account for potential batch effect. For percentage of transfer, we estimated a mixed effect logistic regression model adjusted on the condition, the day and the number of organoids. Second order interactions among condition and day and among number of organoids and day were added to the model in order to normalize statistical tests on time-varying heterogeneity of the number of organoids. A random effect corresponding to replication of the experiment was also added to the model in order to account for potential batch effect. All statistical tests to compare groups (among either cell lines, timepoints or treatments) were deduced by computing contrasts of one the abovementioned logistic model. P-values were therefore adjusted using Tukey’s method. To compare the gene expression measured by RT-qPCR, Holm-Sidak method was applied to determine statistical significance, with alpha=5. ANOVA two-way test was performed to compared cell number at different timepoints of the adherent culture. For the comparison of cell number in tumor organoids, the number of cells was transformed in logarithmic scale and slopes were compared.

## Results

### 1) Patient-derived GBM cells with stem-like features form TNT-like connections

We obtained two GBM cells from a single tumor from a patient in the frame of the clinical trial STEMRI (Identifier: NCT01872221). This trial was aimed at studying the tumoral cells surrounding the core tumor after the latter has been surgically removed, and better understanding and possibly anticipating which of them are at the origin of the relapse. Two bulks of tissue were resected from the infiltrative tumor area defined by the Fluid-attenuated inversion recovery (FLAIR) sequence on MRI (Fig. 1A) and characterized by multimodal MRI spectroscopy for their metabolic activity through the measurement of the Choline/N-AcetylAspartate Index (CNI) [39,40] (Supplementary Fig. 1A). The tissues were then desegregated and cultured in stem cell medium[7] to enrich in GBM stem cells rather than differentiated ones[33], and grown in suspension in neurosphere-like aggregates (Fig. 1B). To further characterize the two populations, named C1 and C2, we monitored the expression of genes associated with different cell types, from differentiated to progenitor/stem cells[4,6,41–44]. GFAP and CHI3L1 (respectively astrocytic and mesenchymal markers) were not expressed and low expression was observed for the neural markers Tubβ3 and GAP43. On the other hand, expression of the progenitor and stem cell markers Olig1, Olig2, Sox11 and Sox2 was significant (Fig. 1C). This pattern of gene expression was retained over passages in culture indicating maintenance of the stemness properties distinctive of GBM stem-like cells (GSLCs) [3]. We further characterized the C1 and C2 cells by Agilent Seahorse XF Cell Mito Stress Test allowing us to monitor mitochondrial function by directly measuring the aerobic respiration of cells. We observed that C2 and C1 cells behave in accordance to their respective region of origin (Supplementary Fig. 1A). Indeed, the C2 population, derived from the highest metabolic area of the external tumoral region (CNI>2), displayed higher maximal respiration and spare respiratory capacity compared to C1 cells originated from a lower metabolically active area (CNI<2). On the contrary, C1 cells presented higher levels of non-mitochondrial oxygen consumption and basal respiratory capacity than C2 cells (n=3, Supplementary Fig. 1B). These data are consistent with a higher mitochondrial oxidative phosphorylation phenotype in the C2 than in C1.

**Figure 1.**
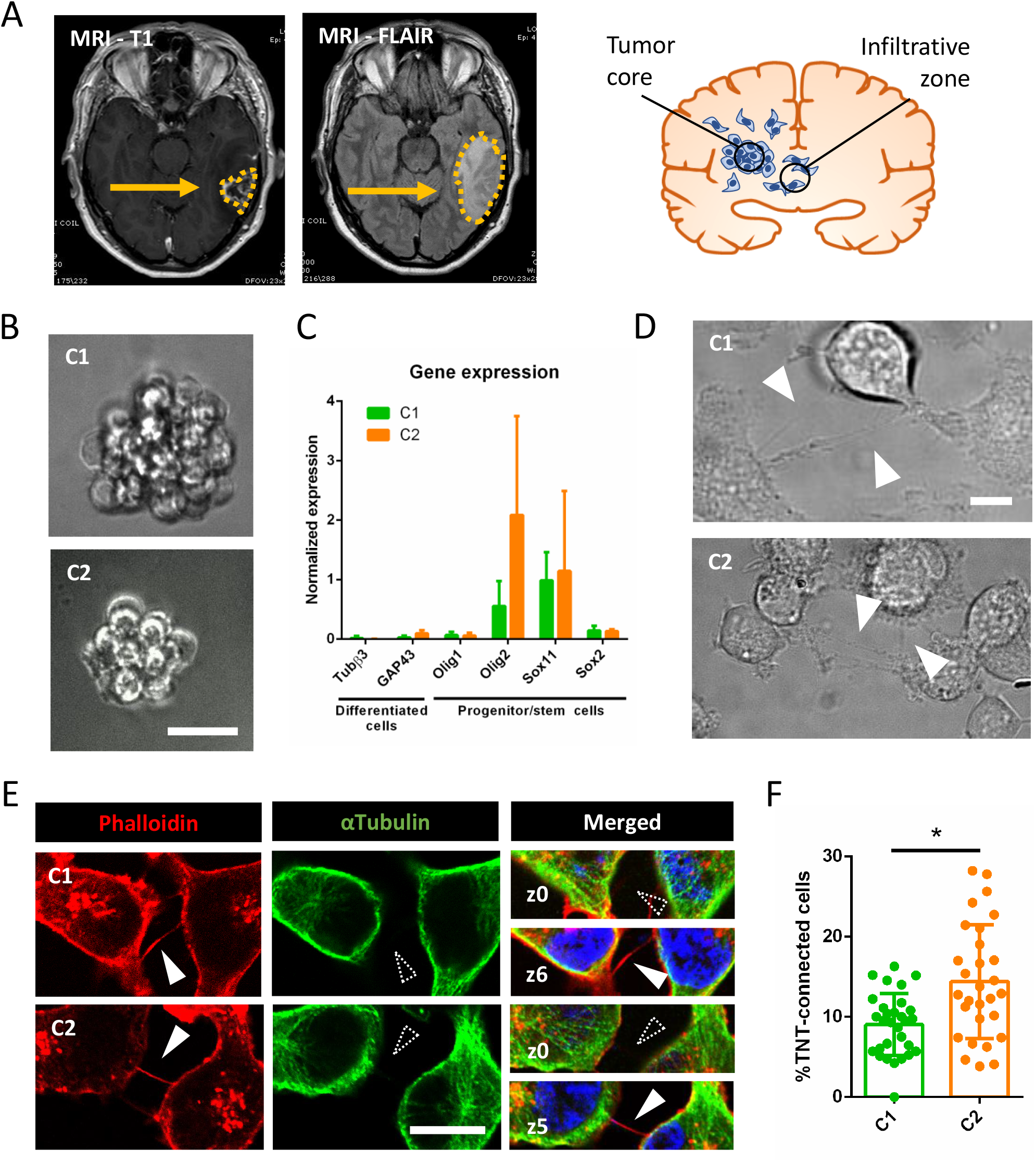
GSLCs form TNT-like structures. (A) MRI analysis of C patient glioblastoma. The tumor is composed by a compact cellular part defined ‘Tumor core’, identified by MRI T1-Gadolinium (on the left). Some tumoral cells infiltrate the normal tissue, forming the ‘Infiltrative zone’ which is identified by MRI-FLAIR (right picture and schematics). C1 and C2 cells were obtained from different parts of the infiltrative zone (see Supplementary Figure 1). (B) C1 and C2 cell growth, forming neurosphere-like clusters in suspension. The resulting images represents a Z-projection of 30 and 50 slides (step size: 0.5 μm), respectively, acquired in Bright field using 40X magnification. Scale bar = 10 μm. (C) Expression of differentiation and progenitor/stem cells markers in C1 and C2, respectively in green and orange. The relative gene expressions were quantified by RT-qPCR after RNA extraction. Data were normalized over the expression of HPRT, housekeeping gene. GFAP and CHI3L1 showed no expression in both C1 and C2 and are not represented on the graph. The graph represents the means with SD of 5 independent experiments, each point performed in triplicate. P values > 0.05 are not significant and not indicated on the figure. (D) GSLCs connected by TNT in live imaging in 2D culture. Cells were seeded on laminin-coated plates and pictures were taken after 6h of seeding using 60 × 1.4NA CSU oil immersion objective lens using Bright field. Arrowheads point to TNT-like connections. Scale bar = 20 μm. (E) GSLC TNTs containing actin but not microtubules. Cells were plated on laminin-coated surface for 6 hours, fixed and stained with phalloidin (actin filaments, red), anti-αTubulin (microtubules, green) and DAPI (nuclei, blue). Representative images were acquired showing TNTs, actin-positive and αTubulin-devoid, floating above the dish surface. White-filled arrowhead indicates presence of TNT labelling, dashed arrowhead indicated absence of TNT staining. Scale bar = 10 μm. (F) Quantification of TNT-connected cells in C1 and C2, respectively in green and orange. GSLC were plated on laminin-coated surface, fixed after 6h and stained with WGA. 2×2 tiles images were acquired with 60x objective and analysed by Icy software. C1 were forming 9.0±4% of connecting cells (5 independent experiments each performed in duplicates, total n cells counted=1239), while C2 were forming 14.4±7% (4 independent experiments, each performed in duplicates, total n cells counted=1367), significantly more connected cells than C1 (p=0.0370). Each dot represents an image containing an average of 40 cells each. P-values were deduced from contrast comparing the two cell populations in a logistic regression model. Error bar = standard deviation. P value < 0.05 (*).

Different GBM-derived cell lines have been described to form TNT-like connections and to be able to transfer cellular content including mitochondria[28–31]. Because such cell lines are only partially recapitulative of the original tumoral features[45], here we aimed to address whether patient-derived GSLCs can form functional TNTs. Thus, C1 and C2 cells were plated for 6h on laminin-coated surface in 2D culture to make them adhere to the support for ease of TNT recognition. Thin cell connections were detected after 6 hours of culture by live imaging (Fig. 1D). After fixation, we assessed the presence of actin-containing connections floating above the laminin coated surface, which is a distinguishing characteristic of TNTs, that hoover above the substrate, as exemplified in Fig. 1E, where both attached-to-the-substrate (z-stack=0) and above (z-stack>3) stacks are shown[35,46]. GSLC TNTs resulted to be always positive for actin and negative for microtubules markers, consistent with the description of classical TNTs[16] (Fig. 1E). For quantification of TNTs, we labelled the cell’s plasma membrane with fluorescent-Wheat Germ Agglutinin (WGA) and counted thin, continuous, non-attached to the substratum protrusions[35] connecting distant cells. Both C1 and C2 populations formed TNT-like connections with a significantly different frequency: about 10% of TNT-connected cells in C1 and 15% in C2 (Fig. 1F). These data showed for the first time that GSLCs, derived from the infiltrative region of the tumor, can interconnect through TNT-like structures.

### 2) TNT-like structures of GSLCs can transfer mitochondria

TNTs are described to be open-ended connections allowing the passage of cellular cargoes. To determine whether the connections observed were apt to this purpose, we decided to assess the transfer of mitochondria, shown to occur in several types of cancer cells (mesothelioma, leukemias, ovarian, etc.)[13,14,27]. To observe mitochondria in living samples, we introduced a GFP-tagged fragment of the subunit VIII of human cytochrome C oxidase located in the inner mitochondrial membrane (MitoGFP) in both GSLCs by lentiviral transduction. We then performed live-imaging on GSLCs and found mitochondria moving inside TNT-like structures and entering into a connected cell (Fig. 2A, Supplementary video 1), supporting an open-ended TNT relying the two cells. To quantify this transfer, we performed co-culture assays[35] between a donor population, expressing MitoGFP, and an acceptor cell population transduced with lentivirus governing the expression of cytosolic mCherry (Fig. 2B). More than 80% of each cell population was stably expressing the constructs, allowing a 1:1 co-culture ratio between the two populations. Cells were plated on laminin-coated surface and grown either in direct contact or separated by a filter (transwell) (Fig. 2B), and the percentage of mitochondria transfer was assessed after 2 or 5 days of co-culture by flow cytometry[35]. Between 1 and 3% of acceptor cells received donor-derived mitochondria, exclusively due to a contact-dependent mechanism since negligible transfer was observed when the two cell populations were separated by filter but shared the same medium (Fig. 2C). Furthermore, the percentage of acceptor cells receiving mitochondria was increasing over time in both GSLCs (Fig. 2D). It is worth noting that C2 had higher mitochondria transfer compared to C1, in accordance with the higher percentage of TNT-connected cells in this population (Fig. 1F). As both GSLCs had a similar proliferation rate in this condition (Fig. 2E) we could rule out that the different transfer abilities of the two cell types was related to differences in cell densities. Next, to confirm that the fluorescence signal detected by flow cytometry corresponded to true mitochondria inside acceptor cells, confocal microscopy was performed in the same co-culture conditions used for the flow cytometry experiments. By this mean MitoGFP puncta (Fig. 2F) which overlapped with TOM20 (Translocase of the Outer Membrane, Supplementary Fig. 2) were observed in acceptor cells. These data indicate that both C1 and C2 GSLCs are able to form functional TNTs when cultured in 2D. However, C1 and C2 transfer mitochondria with distinct efficiencies, consistent with their distinct abilities to form TNTs (Fig. 1F).

**Figure 2.**
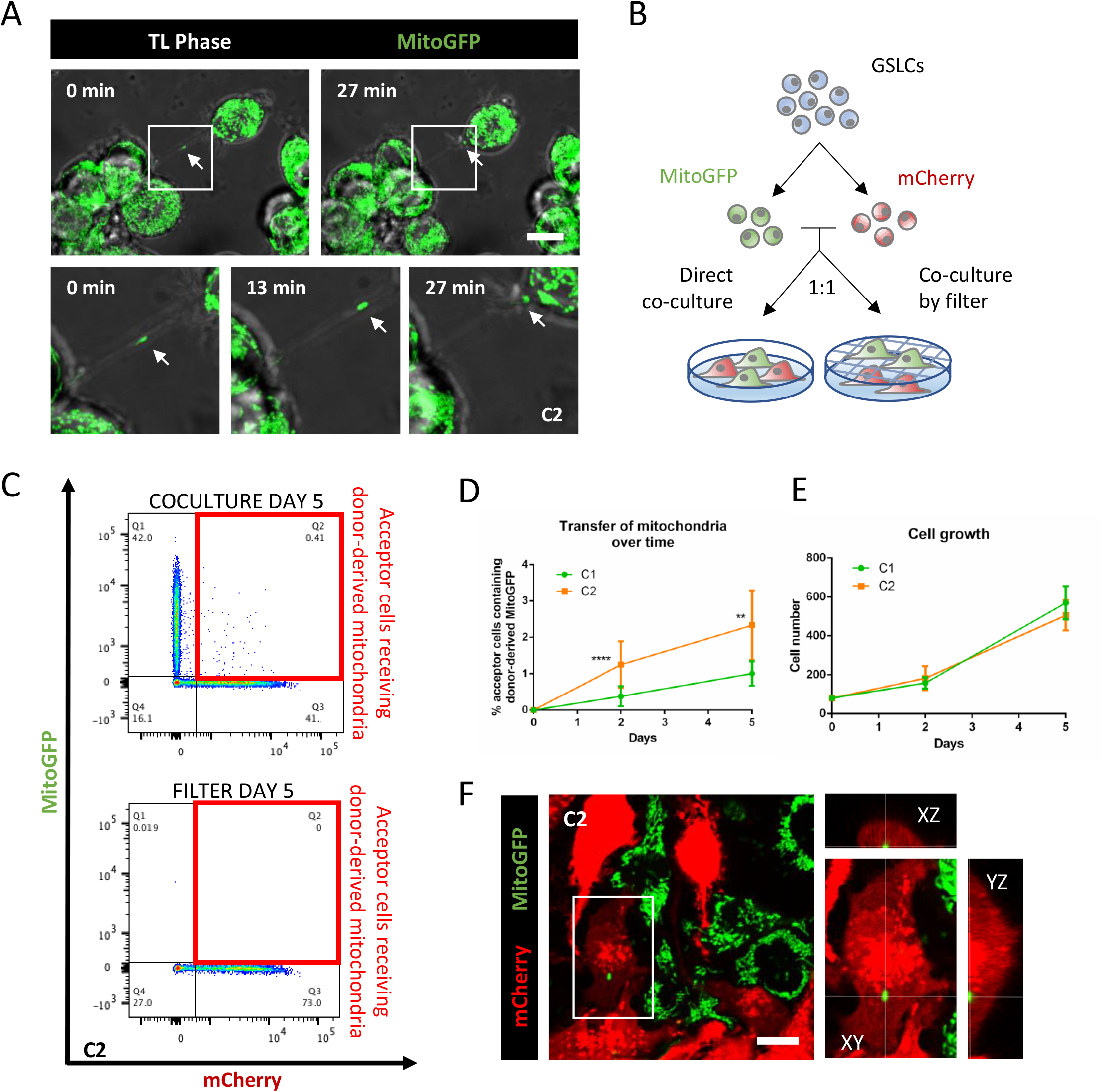
GSLCs transfer mitochondria through TNTs. (A) C2 expressing MitoGFP are connected by TNTs containing mitochondria. Cells were seeded on laminin-coated dish and after 6 hours video were acquired using Bright field and laser 488 in a Spinning Disk microscope. Timeframes show the mitochondria moving along the connection and entering in one of the two connected cells. Each timeframe of the video is the result of the Z-projection of 18 slides (step size: 0.5 μm) (B) Schematic representation of the co-culture experiment. Donor MitoGFP cells were co-cultured with acceptor mCherry cells at 1:1 ratio either by direct contact or through a 1 μm filter. (C) Representative flow cytometry plot of C2 after 5 days of coculture. Acceptor and donor cells respectively lie on the X and Y axis. Acceptor cells positive for MitoGFP signal are framed in the red boxes. (D) Quantification by flow cytometry of the mitochondria transfer over time in C1 and C2, respectively in green and orange. A minimum of 10000 events were analyzed after 2 or 5 days of coculture. C1 shows 0.38±0.27% and 1.01±0.33% of acceptor cells receiving mitochondria after 2 and 5 days, respectively (4 independent experiments, each performed in duplicate). C2 shows 1.25±0.63% and 2.33±0.95% of acceptor cells receiving mitochondria after 2 and 5 days, respectively (5 independent experiments, each performed in duplicate), significantly more than C1 (p<0.0001 (****) at day 2, p=0.0085 (**) at day 5). P-values were deduced from contrast comparing the two cell populations in a logistic regression model. Error bar = SD (E) Cell growth in co-culture experiment. 80000 GSLCs per well were plated at time 0 and counted after co-culture. For C1, 158000±28751 and 568866±85332 cells were counted after 2 and 5 days, respectively (3 independent experiments). For C2, 182900±61890 and 505260±77515 cells were counted after 2 and 5 days, respectively (5 independent experiments). Error bar = SD. ANOVA two-way test was performed and showed no significant difference between C1 and C2 at the two timepoint in analyse. (F) Representative image of co-culture assay in C2. Donor MitoGFP (in green) and acceptor mCherry cells (in red) were fixed after 5 days of co-culture, confocal images were acquired with 63x objective. In the magnification, the orthogonal view of an acceptor cell containing donor-derived mitochondria. Scale bar = 10 μm.

### 3) TNT-like structures exist in GSLC tumor organoids together with TM-like protrusions

In order to assess the participation of the TNT-mediated communication to GBM networking in a context more representative of the tumor, we cultured individually each GSLCs in tumor organoids, according to the protocol published by J. Rich and colleagues[34]. Tumor organoids are a relevant, 3-dimensional, culture method which allows long-term growth preserving the stem cell identity of some cells and reconstitutes, to some extent, the morphological and phenotypic heterogeneity of the original tumor[34]. We were able to grow C1 and C2-derived tumor organoids up to more than 23 days of culture. To characterize the transcriptional changes undergone by the cells in this culture system, we quantified the expression of differentiation and progenitor/stem markers in 23-days-old organoids by RT-qPCR. We observed no significant variation in the expression of all tested genes compared to classical culture, except of GAP43 in C2 cells (Fig 3B), that resulted to be 12-fold higher in organoids. This result showed that although most of the cells still expressed progenitor markers, maintaining therefore their multipotency, some could also commit to a more differentiated profile, as it happens *in vivo*.

**Figure 3.**
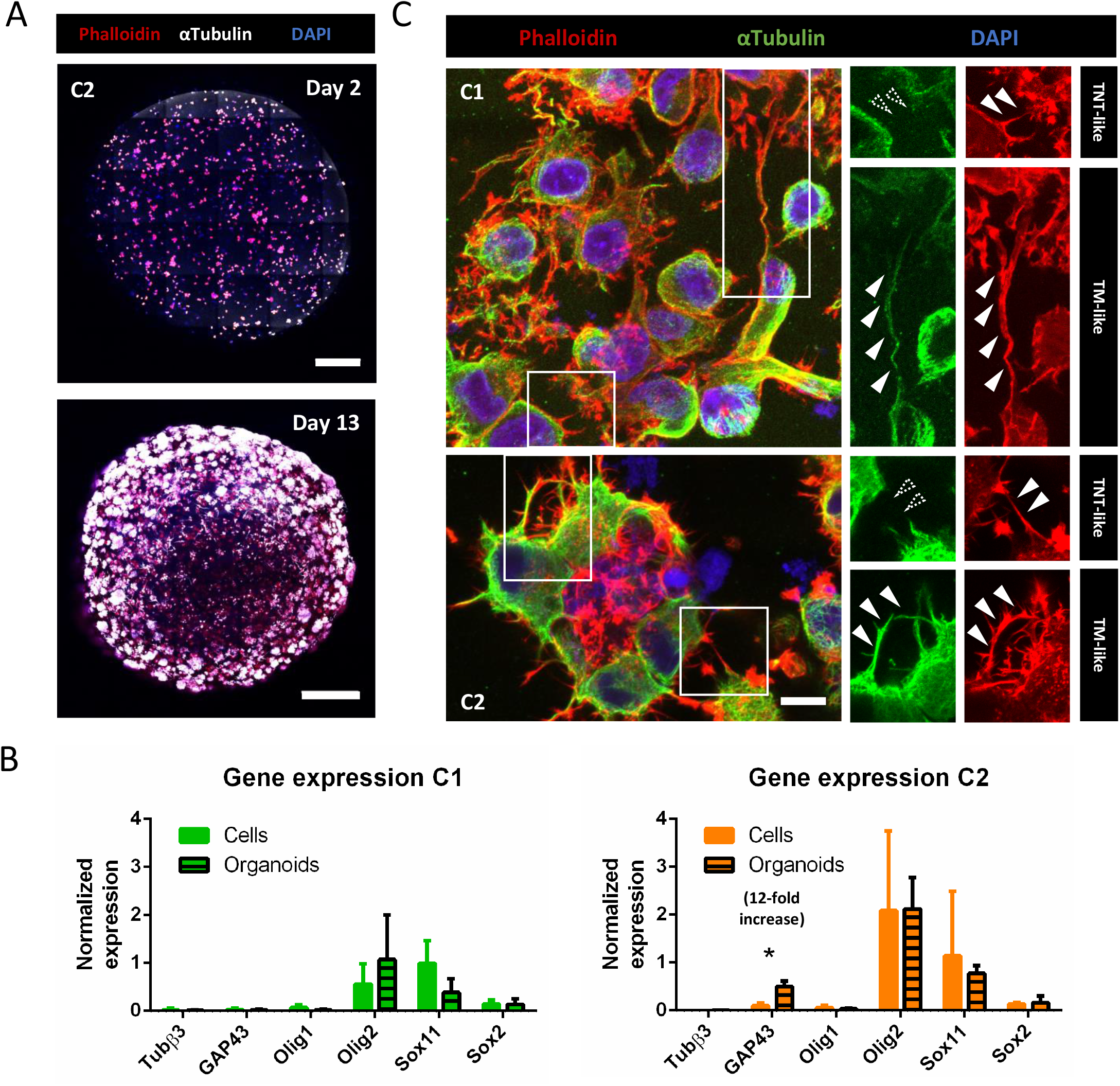
GSLCs in tumor organoids. (A) Image of C2 tumor organoid at 2 and 13 days of growth using Pln-Apo 10X/0.45 objective of inverted confocal LSM700. The resulting images represent a max intensity projection of 5 and 31 sections (step size: 7 and 3.12μm), respectively, stained for anti-αTubulin (microtubules, white), Phalloidin (actin in red) and nuclei (blue). Scale bars are 200 (top) and 500 μm (bottom). (B) Expression of differentiation and progenitor/stem cells markers in C1 and C2 organoids, respectively in green and orange. The relative gene expressions were quantified by RT-qPCR after RNA extraction from 23-days-old organoids, normalized over the expression of HPRT. Note the 12-fold increased expression of GAP43 in C2 tumor organoids, and GFAP and CHI3L1 show no expression in both conditions and are not represented on the graph. The graphs represent means with SD of 3 and 4 independent experiments for C1 and C2 respectively, each point performed in triplicate. Holm-Sidak method was applied to determine statistical significance between cells and organoids for each gene. P value < 0.05 (*), P values > 0.05 are not significant and not indicated on the figure. (C) C1 and C2 tumor organoids at 9 and 6 days, respectively, stained for anti-αTubulin (microtubules, green), Phalloidin (actin filaments, red), and nuclei (blue). Confocal images were acquired with 40X objective. Region of interest show either αTubulin-devoid connections, defined as TNT-like (<1 μm), or thick αTubulin-positive connections (>1 μm), named TM-like. Dashed arrowheads indicate absence of fluorescent signal at the connection level, white-filled arrowhead show positiveness to the signal. Both images are max intensity projections of 12 slices (step size: 0.38 μm). Scale bar = 10 μm.

To date, TNT visualization in 3D cultures and their quantification had not been reported, as preserving and identifying these fragile thin structures in 3D is extremely challenging. In both C1 and C2-derived tumor organoids, fluorescently labelled with phalloidin and anti-αTubulin antibody and imaged with confocal microscopy, we observed different types of cell protrusions (Fig. 3C). Specifically, we noticed thin (<1 μm), actin-rich structures (devoid of tubulin) resembling TNTs that were found connecting cells already within the first week of organoids culture. However, the resolution of light microscopy doesn’t not allow to assess if these connections were functional TNTs or close-ended protrusions (like filopodia) [16]. One possibility to address this, is to look for the presence of organelle inside the connections and assess transfer. To this aim, first we imaged tumor organoids prepared with GSLCs stably expressing MitoGFP construct. Several cell extensions were found to be rich in content of mitochondria (Supplementary Video 2). Of note, by using live-imaging we observed thin TNT-like connections containing mitochondria trafficking between two connected cells (Fig. 4A, Supplementary Video 2, white arrow, and Supplementary Fig. 3), in accordance with what was observed in 2D. Next, to quantify mitochondrial transfer, we prepared tumor organoids mixing MitoGFP donor and mCherry acceptor cells in a 1:1 ratio. After 6, 9, 13, 16, 20 and 23 days of co-culture inside the same organoids, duplicates of 3 organoids per condition were desegregated (and combined) in a single cell suspension and analysed by flow cytometry for the presence of MitoGFP into acceptor mCherry-positive population (Supplementary Fig. 5). The percentage of acceptor cells receiving mitochondria was increasing over time, reaching around 3% in C1 and 8% in C2 after 23 days of culture (Fig. 4B, note the logarithmic y axis scale). Higher efficiency of transfer was observed in C2 cells when comparing the general trend of the transfer with the one of C1 (Fig. 4B), in agreement with the data obtained when cells were cultured in 2D. Mitochondria transfer was not related to cell proliferation as both GSLCs grow similarly in organoids (Fig. 4C). Overall, these data were consistent with the results obtained in 2D and suggested that C2 have higher ability to form and use TNTs for transferring cellular content, compared to C1. To verify that mitochondria transfer was not due to secretion, we co-cultured organoids composed of only one cell population, either donor or acceptor cells, in the same medium. We have not observed any transfer of MitoGFP from donor organoids to acceptor organoids in these conditions over time (Supplementary Fig. 5), strongly suggesting that the mitochondria transfer that we quantified in mixed organoids was dependent on direct cell contacts between donor and acceptor cells. Overall, these data also indicate that GSLCs form functional TNT-like connections able to transfer mitochondria.

**Figure 4.**
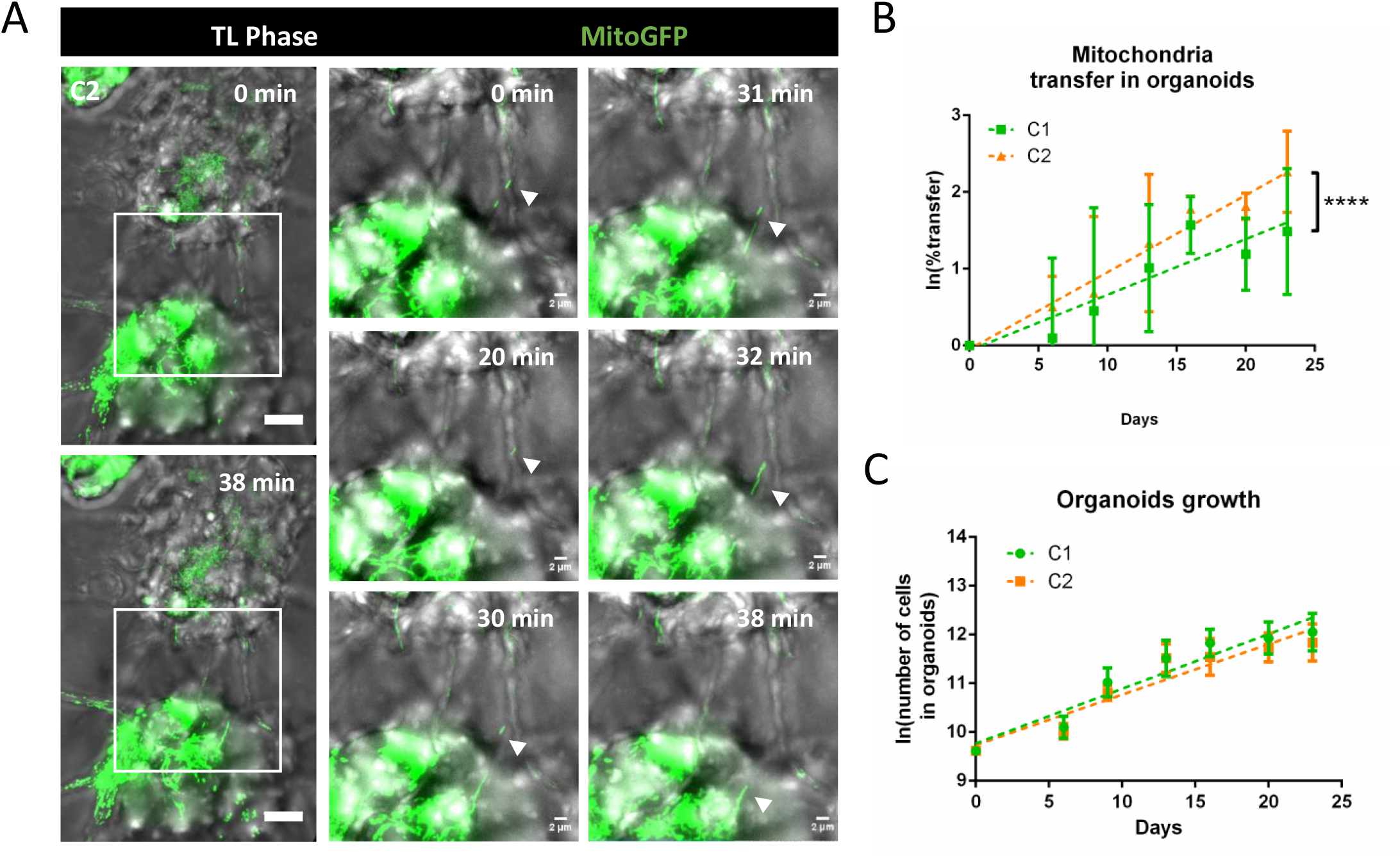
Mitochondria transfer in tumor organoids. (A) TNT-like connection between C2 cells containing mitochondria in 6-days old tumor organoids. Timeframes result of the max projection of 62 slides (step size: 0.5μm) with a total physical thickness of 31μm, with 1 minute of interval time. White arrows point to the mitochondria movement inside the TNT at the different time points. Video were acquired using Bright field and laser 488 in a Spinning Disk microscope. (B) Quantification of the mitochondria transfer in tumor organoids over time in C1 and C2, respectively in green and orange. Organoids were prepared mixing donor and acceptor cells for each GSLC. For each timepoint and condition, duplicates of a pool of 3 organoids were dissociated in a single cell suspension and fixed for flow cytometry analysis after 6, 9, 13, 16, 20 and 23 days of culture. All the cells in the suspension were analyzed to obtain the percentage of acceptor cells receiving mitochondria. C1: day 6 1.54±1.4%; day 9 2.80±2.9%; day 13 2.20±1.1%; day 16 5.07±2.06%; day 20 3.55±1.5%; day 23 3.05±0.84% (4 independent experiments). C2: day 6 1.72±0.7%; day 9 2.64±2.2%; day 13 4.96±4.35%; day 16 5.98±1.02%; day 20 5.57±0.03%; day 23 8.37±2.7% (3 independent experiments). Percentage of transfer was transformed into a logarithmic scale. Error bar = SD. P-values are deduced by comparing the slopes of the two cellular population in a logistic regression model as described in material and methods. P value < 0.0001 (****) (C) Cell number in tumor organoids. For each timepoint and condition, duplicates of a pool of 3 organoids were dissociated in a single cell suspension C1: day 6 24800±5768; day 9 63150±18350; day 13 105850±43970; day 16 140450±33929; day 20 158600±60394 day 23 181800±78820 (4 independent experiments). C2: day 6 22600±3704; day 9 49700±8116; day 13 104200±33870; day 16 108580±42218; day 20 128800±34478; day 23 145080±47726 (4 independent experiments). The cell number was transformed into a logarithmic scale and slopes were compared by linear regression (dashed lines). No significant difference was observed between C1 and C2.

Interestingly, in addition to the TNT-like connections described above, we also observed thick (>1 μm) and long protrusions containing both actin and microtubules rather similar to TMs (previously observed *in vivo*[10], but not in the 2D cultures), which coexisted with TNTs in the same organoids (Fig. 3C).

To better characterize TM-like protrusions we decided to evaluate the presence of GAP43, a TM specific marker described to be one of the major driver of TM formation [10]. Intriguingly, by RT-qPCR we had observed increased expression of this marker in C2-but not in C1-derived tumor organoids (Fig. 3B). In agreement with these data, by immunofluorescence we observed an increase in the number of GAP43-positive C2 cells over time (see days 2, 6 and 13 labelling’s in Fig. 5A, and B), confirming that tumor organoids reproduced to some extent tumoral heterogeneity and structure. In addition, part, but not all, of TM-like extensions resulted positive for GAP43 (Fig. 5C) in C2 organoids. Of interest, some of these thick TM-like structures in C1 and C2 organoids contained mitochondria (Supplementary video 2, red arrow). This was consistent with the observations of Winkler and colleagues [10], and also in our case mitochondria transfer was not detected through such structures[47]. The presence of these two types of connections, TM-like (also expressing GAP43) and TNT-like, was further confirmed in organoids produced from one GSLCs (O) derived from a second patient (Supplementary Fig. 4), suggesting that the coexistence of TNTs and TMs in GBM could be recurrent in different tumors. Taken together, these results showed that the 3D organoid model using GSLCs is a valid representation of the tumoral complexity *in vivo*[34] and that various types of connections, including TNTs and TMs, could coexist in the network formed by GSLCs. However, only TNTs could provide a route for the exchange of cellular material, notably mitochondria.

**Figure 5.**
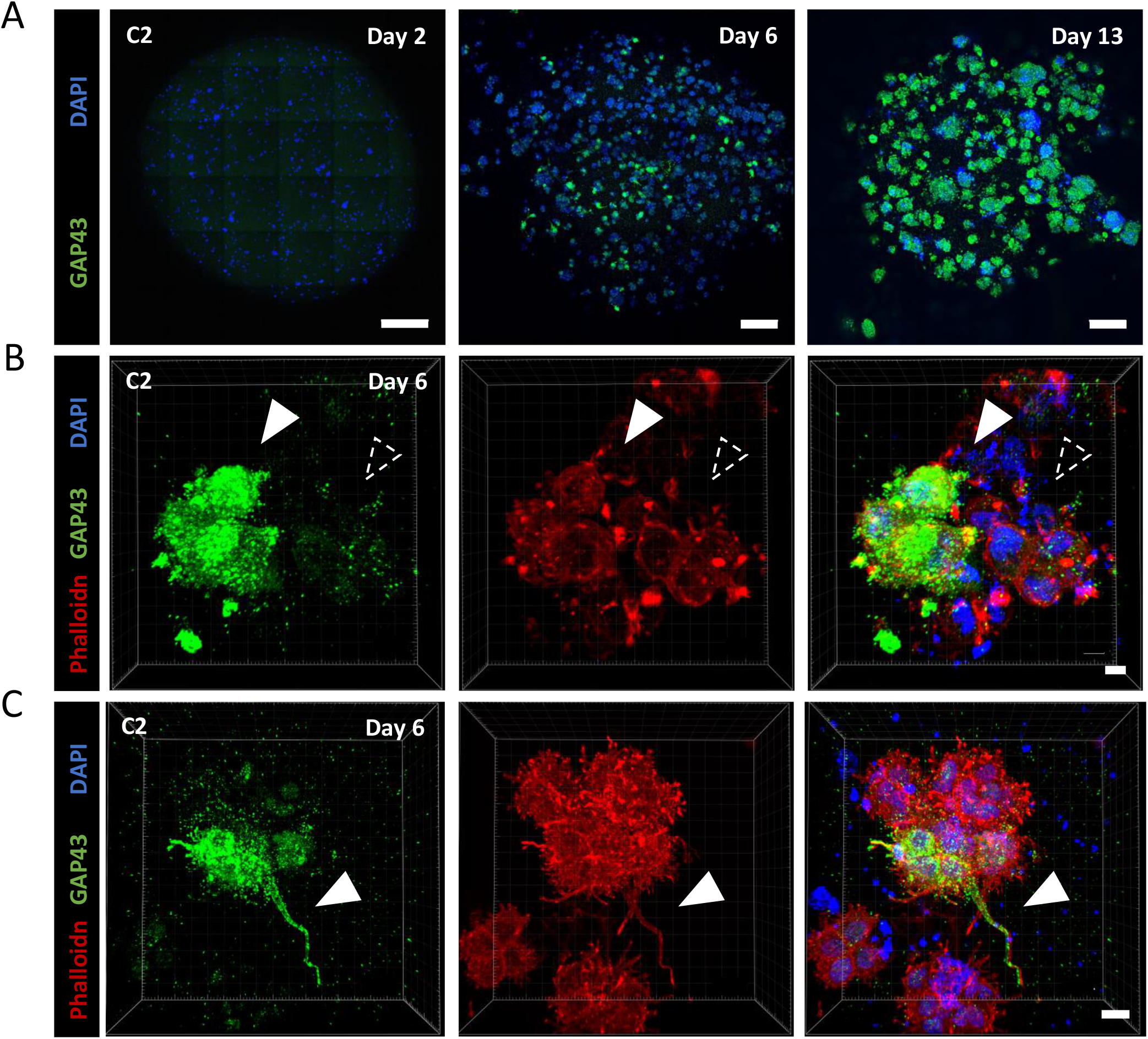
GAP43 expression and TM characterization in tumor organoids of GSLC cells. (A) GAP43 protein expression increases over time. 2, 6 and 13 days-old organoids were fixed and stained with anti-GAP43 (in green) and DAPI (in blue). Confocal images with 10x objective were acquired. Images result from the max intensity projection of 5, 20, 11 sections (step size: 7, 3.13, 3.13 μm), respectively. Scale bars: 500 μm, 200 μm, 200 μm (from left to right). (B) Heterogeneous expression of GAP43 in C2 tumor organoids. 6 days-old C2 organoids were fixed and stained with anti-GAP43 (in green), phalloidin (actin filaments, in red) and DAPI (in blue). Confocal images with 63x objective were acquired. 3D reconstruction of a 50-sections image (step size: 0.33 μm) was performed using Imaris Viewer software. White-filled arrowhead point to a cluster of cells expressing GAP43, alternatively a group of cells negative for its expression are indicated with a dashed arrowhead. Scale bar: 5 μm. (C) TM-like protrusion can express GAP43 in C2 organoids. 6 days-old C2 organoids were fixed and stained with anti-GAP43 (in green), phalloidin (actin filaments, in red) and DAPI (in blue). Confocal images with 63x objective were acquired. 3D reconstruction of a 77-sections image (step size: 0.77 μm) was perfomed using Imaris Viewer software. White-filled arrowheads point toward a TM-like extension expressing GAP43. Scale bar: 15 μm. 3D reconstructions were performed with Imaris Software.

### 4) Effect of irradiation on TNT-based communication in C1 and C2 GSLCs

GBM relapse, originates from GSLCs [4] and treatments introduce alteration in GBM cells that seem to favors their survival [7]. As TNTs have been involved in cancer progression of different tumors [13] we decided to assess the effect of irradiation on the TNT-based communication in GSLCs. We irradiated cells at a dose of 2 Gray (Gy). This dose, daily administrated to GBM patients for five days a week for six weeks during radiotherapy [1], was applied only once on our cells in order to be effective but subtoxic[7] and preserve cell viability for all the duration of the experiment. Cells were plated on laminin-coated coverslips for 6 hours at 1, 3 and 6 days after the irradiation, and next were fixed and analyzed for their TNT content. While C1 cells showed a slight decrease, not statistically significant, in their percentage of TNT-connected cells after irradiation, TNT frequency was significantly increased in C2 cells the day following the irradiation (Fig. 6A), suggesting an acute effect induced by the irradiation in this population. To assess the effect of irradiation on the transfer of mitochondria, 2 Gy irradiation was applied on the donor cells the day before the co-culture and transfer was quantified after 2 or 5 days of co-culture. The percentage of C1 acceptor cells containing donor-derived mitochondria was not affected by irradiation, whereas we observed a tendency to an increased transfer upon irradiation in C2 (Fig. 6B), although not statistically significant. Of note, in this experiment, C1 cells showed mild but significant reduction in their cell number at 5 days from irradiation, compared the control condition, differently from C2 cells which did not show significative variations (Supplementary Fig. 6A). To assess variations of mitochondria, transfer upon irradiation on a long-term co-culture we irradiated C1 and C2-derived organoids at day 5 of culture and assessed mitochondria transfer at various timepoints. After 23 days in culture, 3% and 1.7% of transfer was observed, respectively, in C1 control and irradiated condition, showing a significant decrease of transfer in the irradiated condition (Fig. 5C). On the other hand, in C2 there was no significant reduction of the transfer efficiency over time, with 8% and 7% of transfer observed in control and irradiated condition, respectively (Fig. 6C). The different behaviour of C1 and C2 cells was independent of the effect of irradiation on cell growth as the number of cells into both types of organoids was not significantly affected by irradiation (Supplementary Fig. 6B), as expected with the chosen doses. Importantly, when co-culturing single population organoids in the same dish to assess the contribution of secretion in transfer we did not observe transfer.

**Figure 6.**
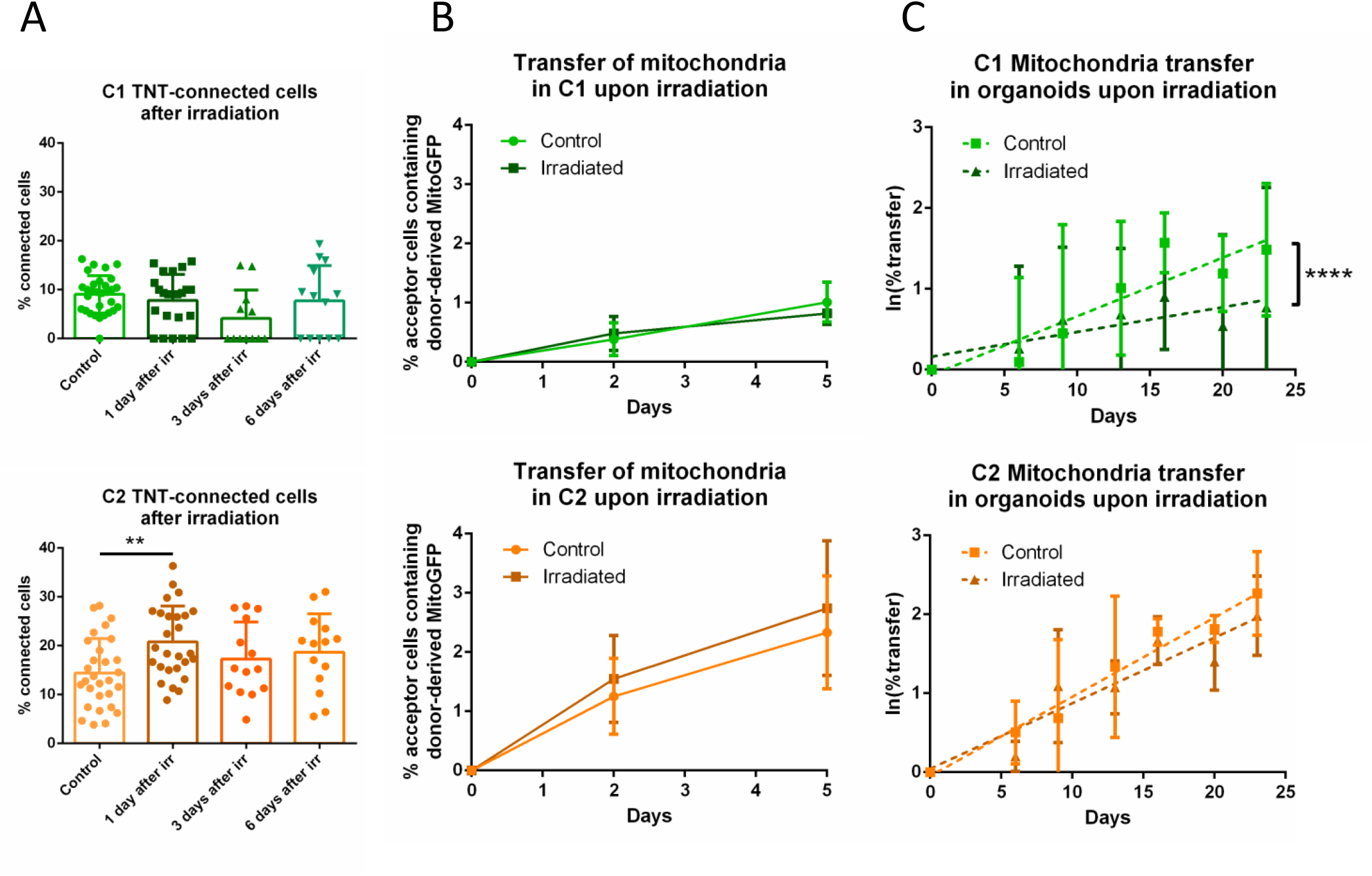
Effect of GSLC irradiation on TNT-based communication. (A) Quantification of TNT-connected cells in C1 and C2 after irradiation, respectively in green and orange, in adherent cell culture. GSLCs were irradiated 1 day before cell plating on laminin-coated surface, then fixed after 6h and stained with WGA. 2×2 tiles images were acquired with 60x objective and analysed by Icy software, experimental duplicates were performed for each condition. The graphs represent means with SD. C1 were forming 7.8±5% (4 independent experiments, tot n cells counted=891), 4.1±6% (3 independent experiments, total n cells counted=300) and 7.7±7% (3 independent experiments, total n cells counted=313) of connecting cells after 1, 3 and 6 days from the irradiation, respectively. No statistically significant difference was observed compared to control (9.0±4%, 5 independent experiments, total n cells counted=1239). C2 were forming 20.8±7% (4 independent experiments, total n cells counted=1368), 17.3±7% (3 independent experiments, total n cell counted=552) and 18.7±8% (3 independent experiments, total n cells counted=462) of connecting cells after 1, 3 and 6 days from the irradiation, respectively. A statistically significant increase was observed 1 day after irradiation compared to control (14.4±7%, 4 independent experiments, total n cells counted=1367, p=0.0073 (**)). Each dot represents an image containing an average of 40 cells each. P-values were deduced from contrast comparing the two cell populations in a logistic regression model. (B) Quantification of the mitochondria transfer by flow cytometry in both GSLCs over time in C1 and C2 upon irradiation, respectively in green and orange. Donor GSLC were irradiated 1 day before the coculture, analysis was performed after 2 or 5 days (corresponding at 3 and 6 days from the irradiation). A minimum of 10000 events were analyzed per condition, each performed in duplicate. In irradiated condition, C1 show 0.48±0.28% and 0.81±0.18% of acceptor cells receiving mitochondria after 2 and 5 days, respectively (4 independent experiments). No statistically significant difference was observed compared to control (day 2: 0.38±0.27%; day 5 1.01±0.33%; 4 independent experiments). In irradiated condition, C2 show 1.54±0.73% and 2.74±1.13% of acceptor cells receiving mitochondria after 2 and 5 days, respectively (5 independent experiments). No statistically significant difference was observed compared to control (day 2: 1.25±0.63%; day 5: 2.33±0.95%. 5 independent experiments). P-values were deduced from contrast comparing the two cell populations in a logistic regression model. Graphs are means with SD. (C) Quantification of the mitochondria transfer in tumor organoids upon irradiation in C1 and C2, respectively in green and orange. Organoids were prepared mixing donor and acceptor cells for each GSLC and irradiated at 5 days from their preparation. Experiment was performed as in Fig 4D. Control C1: day 6 1.54±1.4%; day 9 2.80±2.9%; day 13 2.20±1.1%; day 16 5.07±2.06%; day 20 3.55±1.5%; day 23 3.05±0.84%. Irradiated C1: day 6 1.90±1.6%; day 9 4.45±1.9%; day 13 2.50±1.7%; day 16 2.82±1.5%; day 20 2.39±1.61%; day 23 1.76±1.2% (4 independent experiments, p<0.0001, ****). Control C2: day 6 1.72±0.7%; day 9 2.64±2.2%; day 13 4.96±4.35%; day 16 5.98±1.02%; day 20 5.57±0.03%; day 23 8.37±2.7%. Irradiated C2: day 6 1.23±0.2%; day 9 3.50±2.3%; day 13 3.03±0.9%; day 16 5.46±1.5%; day 20 4.23±1.3%; day 23 7.21±1.7% (3 independent experiments, p = 0.0665). Percentage of transfer was transformed into a logarithmic scale. P-values are deduced by comparing the slopes of the two cellular population in a logistic regression model as described in material and methods. Error bar = SD.

Overall, consistent with the wide heterogeneity present in GBM tumors where distinct molecular profiles coexist and exhibit differential therapeutic responses[3], our data indicated that GSLCs derived from the same tumor display intrinsically different properties and have diverse response to irradiation regarding the effect on TNT formation and transfer function. Specifically, TNT functionality was preserved in C2 organoids after irradiation whereas it was reduced in C1 organoids.

## Discussion

Tunneling nanotubes (TNTs) are gaining an increasing relevance in the context of cancer development and progression[13]. Their presence and ability to transfer cellular material including organelles, has been correlated with the induction of migratory ability, angiogenesis, cell proliferation ad therapy-resistance[14,27]. Few reports have addressed the presence of TNTs in GBM, where it was shown that GBM-derived cell lines in culture were able to form TNT-like connections [28,29,32] and exchange mitochondria with healthy cells of the tumor microenvironment, while promoting cell aggressive phenotypes as increasing proliferation and treatment-resistance [30,31,48]. Nevertheless, at transcriptomic level GBM cells lines are drastically divergent from the original tumor [45], thus they do not appear to be a very relevant model for GBM. On the other hand, TNTs were not reported in patient-derived GSCs xenografted into murine brains, which formed a complex tumor cell network based on thicker neurite-like connections, called TMs [10]. In this elegant work the authors used live two-photon microscopy to image the implanted tumor with a resolution that might have not been sufficient to visualize TNTs. Thus, whether TNTs participate to this networking in a tumoral relevant model, but were not detected because of the approach used, is not yet clear. To address this question, here we have assessed their presence and functionality in two GSLCs derived from the most external tumoral area, responsible for GBM relapse. By using live-imaging and quantification of transferred mitochondria, we showed for the first time that GSLCs are able to interconnect and transfer mitochondria *via* TNT-like structures in 2D culture, in both GSLCs populations. Importantly, to visualize and characterize these connections and to monitor their ability to transfer mitochondria in a context closer to the tumor, we cultured these GSLCs as tumor organoids, a very relevant model shown before to recapitulate at phenotypic and transcriptomic level patient tumors heterogeneity[34,45,49]. Indeed, in our organoid cultures, GSLCs maintained their progenitor characters over more than 23 days. Using confocal microscopy in fixed conditions, we provided evidences for the existence of thin TNT-like protrusions connecting different cells in the organoids from early time in culture; while live-imaging demonstrated mitochondria moving along TNTs and entering into the connected cells. These observations supported the hypothesis that TNTs exist and are functional in GMB derived tumor organoids, suggesting that TNT-based communication may be relevant in the actual tumor. Because TNT communicative abilities were observed in the GSLCs obtained from the infiltrative and putative relapse-driving area, we could speculate that this might have a potentially relevant role in tumor recurrence and induction of aggressive phenotypes, as shown in the case of other tumors [13,14]. Nevertheless, the aggressiveness of GBMs has been correlated before with the establishment of tumor networking based on TMs [10]. TMs are neurite-like extensions[50], able to connect to distant cells and propagate action potential[51,52] and proposed to scaffold the network formed by cancer cells. Interestingly, we also observed thicker connections resembling TMs in our GSLC tumor organoids. Although TMs are organelle-rich structures, no cargo transfer through them has been demonstrated[47], while the presence of GAP-junctions along their length[53] support their role in electric signal transmission. Our data confirmed these observations in organoids, since we could follow movements of mitochondria in cell bodies and protrusions, including TM-like structures. However, the effective transfer of mitochondria until entering a connected cell could be observed only through TNT-like connections, and not in TM[10,11,51]. Of note, we observed and quantified mitochondria transfer in both C1 and C2 GSLCs, grown in 2D culture conditions where none of them expressed GAP43, which is a typical marker and the major known driver of TMs[10,11], and in 3D organoids where only C2 subpopulations expressed GAP43, suggesting that the transfer of mitochondria does not occur through GAP43-dependent structures. Notwithstanding, we observed that C2 cells, which can express GAP43, were transferring mitochondria more efficiently compared to C1 cells, even after irradiation. These results are in accordance with the work of F. Winkler and collaborators[10,11] suggesting that GAP43 expression and the presence of TMs could be correlated to the aggressiveness of the tumor. Because we observed TNTs and TMs in the same organoid, we propose that TNTs and TMs coexist and cooperate in GBM networking, carrying on complementary roles that could participate eventually to treatment-resistance. In particular, we hypothesize that in a situation in which a TM tumor network is formed based on signalling exchanges between interconnected cells there will be the possibility to form more TNTs, which will provide the transfer function and material exchange (Fig. 7). This hypothesis needs to be further tested *in vivo*, ideally in the conditions in which TMs have been observed before [10].

**Figure 7.**
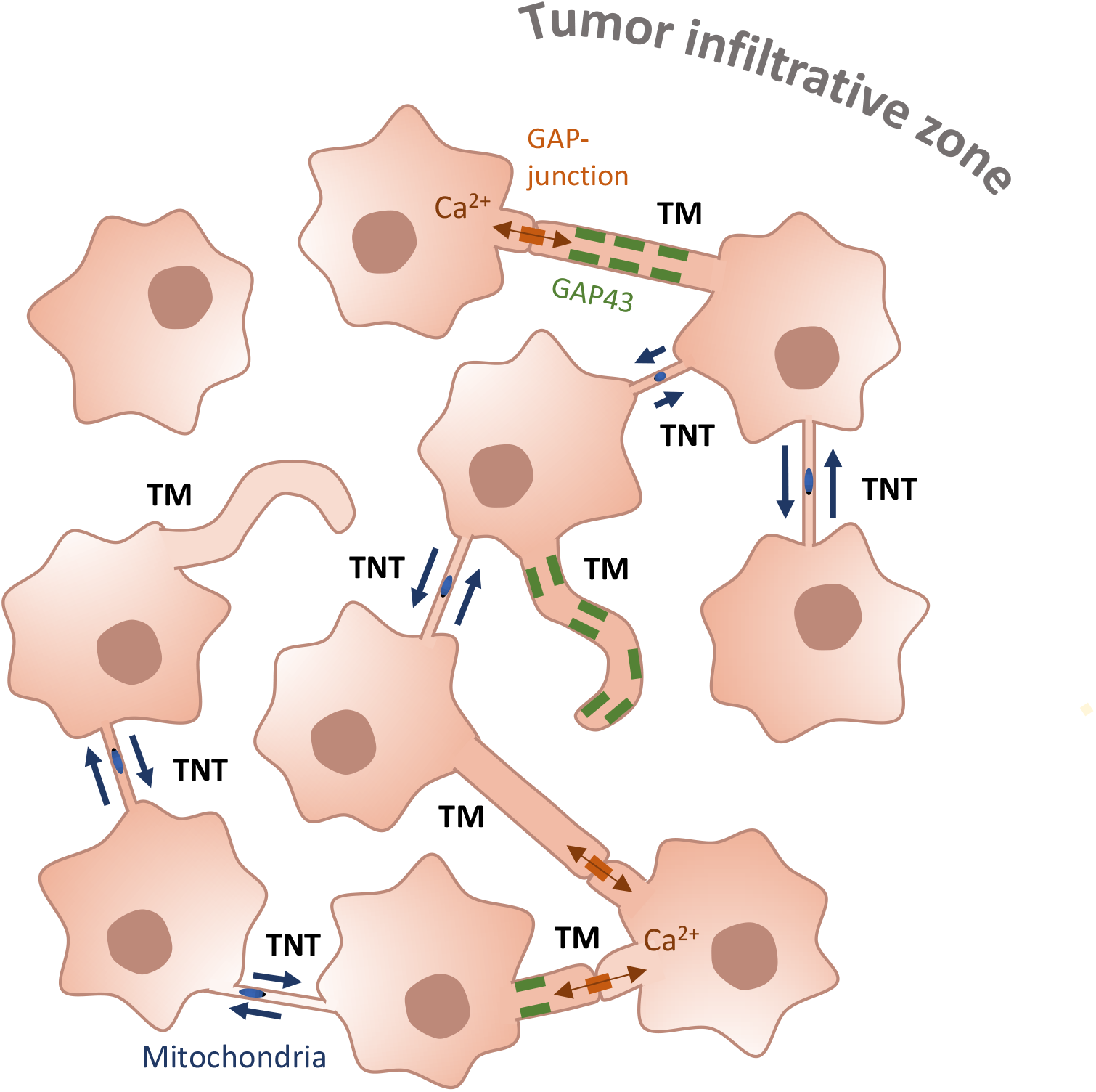
GSLC network. GSLCs interconnect through different types of cellular extensions. TMs are thick (>1μm) protrusions that can either contact other cells through GAP-junctions, allowing the propagation of calcium flux, or be individual finger-like extensions not connecting remote cells. They can be positive for GAP43 (rectangles along the membranes of TM), neuronal Growth-Associated Protein. GSLCs also interconnect through TNTs, thinner (<1μm), open-ended connections which allow transfer of cellular cargos, such as mitochondria (ovals in TNTs).

In our GSLC model we have chosen to specifically look at mitochondria transfer; as mitochondria can provide metabolic support to cancer cells[14,27], transfer of mitochondria has been shown to modulate the response to treatments in a beneficial manner for the recipient cells impacting on their metabolism, rescuing their aerobic respiration and providing a metabolic support against treatment-related stress[27,54]. However the observation of mitochondria transfer between GSLCs does not preclude the possibility that other cellular material (e.g. RNA, proteins, other vesicles) could be additionally transferred through the same connections, as observed in other cancer types[55–57]. Because of the limitation of our study, it will be of important in the future to address first what is the exact composition of transferred material, and second what are the functional consequences of such events for both donor and recipient cells.

Wide cellular heterogeneity is one of the many reasons that makes GBM very difficult to treat, as it is formed by cells with different abilities to respond to the treatments. We have shown that two GSLCs derived from two areas of the same tumor, also display different behaviour, including percentage of TNT-connected cells and transfer of mitochondria over time in response to irradiation. Specifically, C2 cells, coming from the most metabolically active area of the tumor (CNI+) previously shown to be more likely leading tumor relapse [58], in our hands also displayed higher ability to respond to an energetic demand (maximal respiration and spare respiratory capacity, Supplementary Fig. 1C) compared with C1 cells. It is conceivable that the presence of metabolic heterogeneity may offer to GBM tumor cells adaptive mechanisms to better respond and overcome the cellular stress introduced by the treatments [59]. As C2 cells were shown to maintain their TNT-communicative ability after irradiation, differently from C1 cells, it is tempting to speculate that TNTs might contribute to this rescue process. However, the potential link between metabolism and ability to grow TNTs and to respond to treatment needs to be directly investigated by functional studies and in a large number of patient-derived cells. Furthermore, our data do not exclude that other mechanisms (including release of various extracellular factors as well as TMs) are involved in GBM therapy-resistance. TNT-mediated communication has been described to be an advantageous feature for tumoral cells in several cancer models [13], the ability to exploit this way of communication could be common in more aggressive cancer cells, also in the case of GBM. Overall, our data are consistent with tumor networking being important in GBM progression, resistance to treatment and relapse. In addition to TMs, the ability to grow functional TNTs could participate to the formation of a functional tumor network, where exchanges of organelles and different material in addition to signalling molecules is allowed (Fig. 7). Although our data represent a step up to the use of GBM derived cell lines, it will be necessary to further investigate their relevance in a larger sample of tumors to specifically analyse the impact of TNT based networking in the context of tumor progression, therapy resistance and relapse. In this context, it may as well be possible to exploit them by considering inhibition of TNT-dependent transfer as a new way for optimization of radiotherapy efficacy in GBM.

### Conclusions

Our data suggest that TNT-mediated exchange of cellular material occurs between GSCs and that in addition to TMs, TNTs participate to GBM tumoral networking providing a route for the transfer of intracellular material and potentially contributing to tumor progression and treatment-resistance.

## Supporting information

Supplementary video 1: Movement of mitochondria along TNT-like connections in 2D-conditions.

Supplementary video 2: Transfer of mitochondria via TNT-like connections in tumor organoids.

Supplementary video 3: Motion of mitochondria inside TNT-like connections in tumor organoids.

## Abbreviation

GBM: Glioblastoma
GSCs: Glioblastoma Stem Cells
GSLCs: Glioblastoma Stem-Like Cells
TNTs: Tunneling Nanotube
TMs: Tumor Microtube
Tubβ3: Class III β-Tubulin
CHI3L3: Chitinase 3-Like-3
GFAP: Glial Fibrillary Acidic Protein
GAP43: Growth-Associated Protein 43
Olig1: Oligodendrocyte transcription factor 1
Olig2: Oligodendrocyte transcription factor 2
Sox2: Sex-determining-region-Y-Box Transcription Factor 2
Sox11: Sex-determining-region-Y-Box Transcription Factor 11
TOM20: Translocase of Outer Membrane
OCR: Oxygen consumption rate
FCCP: Carbonyl cyanide-4-(trifluoromethoxy) phenylhydrazone

## Declarations

### Ethical approval

The consent for the use of human material has been given with the clinical trial STEMRI (Identifier: NCT01872221).

### Consent for publication

The consent for the use of human material has been given with the clinical trial STEMRI (Identifier: NCT01872221).

### Availability of data and materials

The datasets used and/or analysed during the current study are available from the corresponding author on reasonable request.

### Competing interests

The authors declare no conflict of interest

### Fundings

This work was funded by grants from HTE (HTE201502) to CZ and EMJC, Institut National du Cancer (PLBIO18-103) to CZ and EMJC, Fondation de France (WB-2019-19194) to ISSM and Fondation ARC pour la recherche sur le cancer to GP (DOC20190508549).

## Acknowledgements

We thank all the members of the MOGLIMAGING network, HTE program, Aviesan and INSERM which participated in our collaboration. We thank I. Leroux for teaching tumor organoids preparation and culture. We thank all the members of the UTRAF unit for their support.

## Authors’ information

Giulia Pinto^1,2^, Inés Saenz-de-Santa-Maria^1^; Patricia Chastagner^1^, Emeline Perthame^3^, Caroline Delmas^4^, Christine Toulas^4^, Elizabeth Moyal-Jonathan-Cohen^4^, Christel Brou^1^, Chiara Zurzolo^1^.

---

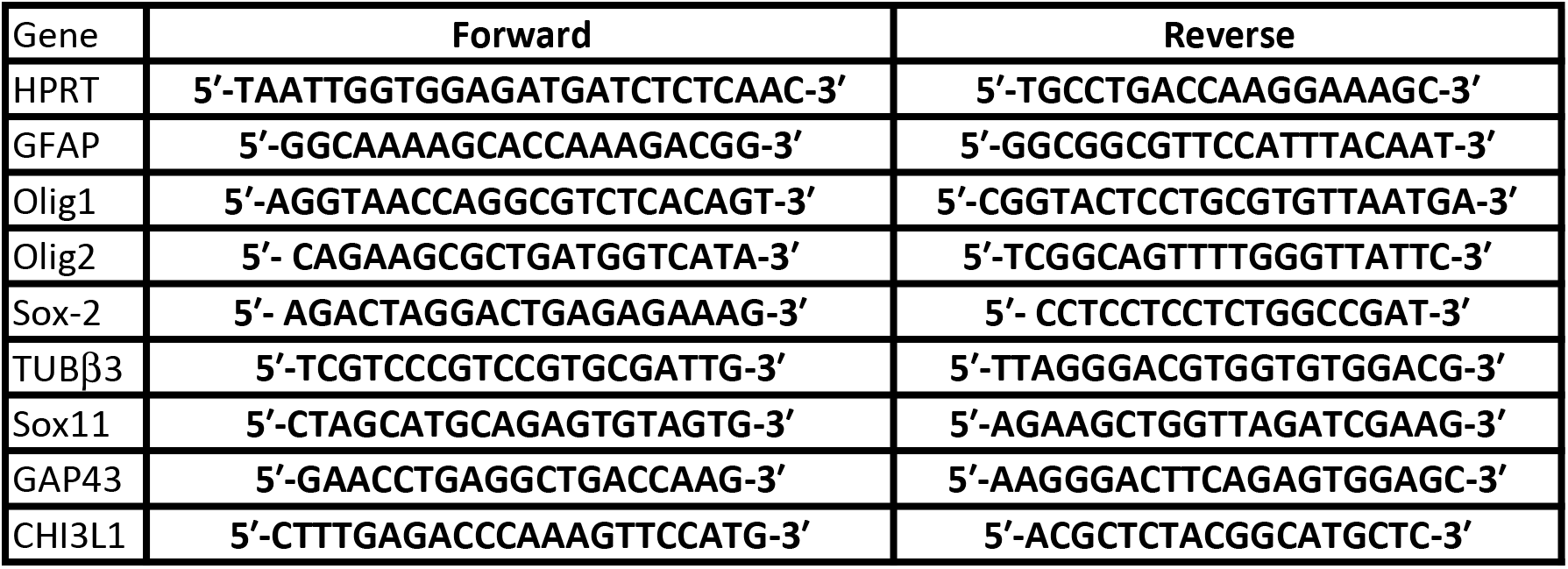

## Supplementary material legends

**Supplementary Figure 1.**
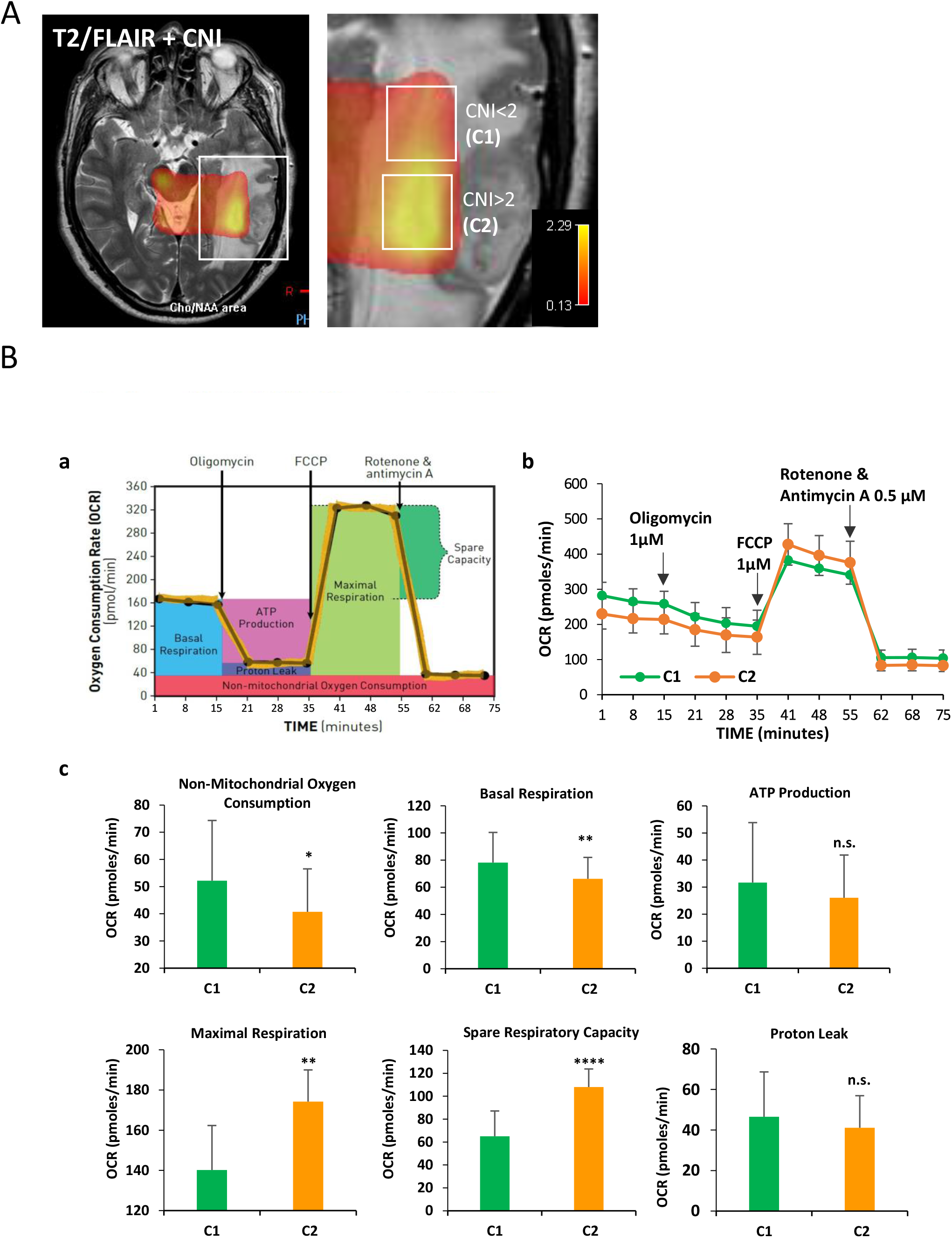
Metabolic features of C1 and C2 cells. (A) MRI-FLAIR imaging of C patient glioblastoma co-registered with functional MRI for the measurement of CNI (Choline/N-AcetylAspartate Index), indicative of the metabolic activity of the tumor [39,40]. The enlargement on the infiltrative area shows the two tumoral regions resected to obtain C1 cells (are with CNI<2) and C2 cells (are with CNI>2). (B) Measurement of mitochondrial aerobic respiration profile using the Extracellular Flux Analyzer Seahorse XF96 **(a)** Schematic of the XF Cell Mito Stress Test used to determine the oxygen consumption rate (OCR) (https://www.agilent.com). **(b)** C1 and C2 cells were seeded in the Seahorse Bioscience microplates laminin-precoated (20,000 cells/well). After 12h, 1 μM oligomycin, 1 μM FCCP (carbonyl cyanide-4-(trifluoromethoxy) phenylhydrazone), and 0.5 μM rotenone/antimycin A were subsequently added. **(c)** Individual parameters for respiration, including non-mitochondrial oxygen consumption, basal respiration, proton leak, maximal respiration, spare respiration capacity, and ATP production, in C1 vs C2 cells. Data are presented as the mean ± standard deviation (3 independent experiments, 10 replicates each condition). Each data point represents an OCR measurement. *p < 0.05, ** p <0.005, **** p <0.00005 C1 vs C2.

**Supplementary Figure 2.**
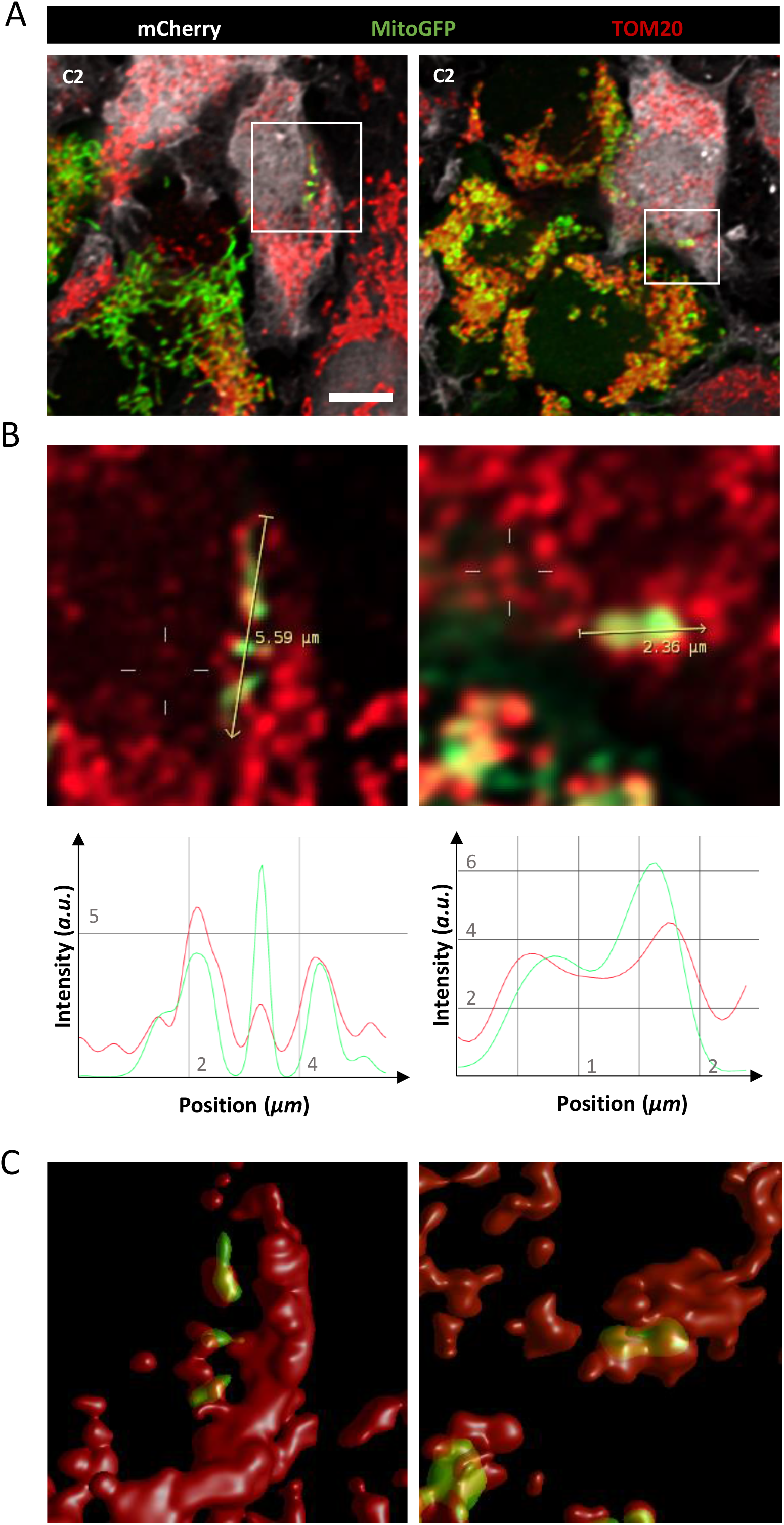
MitoGFP signal in acceptor cells matches with TOM20 mitochondrial marker staining. (A) C2 MitoGFP (in green) and mCherry (in white) cells were co-cultured over 5 days on laminin-coated coverslips, then fixed and stained with anti-TOM20 (in red), mitochondrial marker. Confocal images were acquired with 63x objective and deconvolved with Huygens software. Acceptor cells containing donor-derived MitoGFP signal overlapping wit TOM20 staining were observed (z-stack=2; step size= 0.35 μm), Scale bar 5 μm. (B) A yellow arrow was drawn along the green mitochondria to obtain the intensity profile of MitoGFP and TOM20 signal. The two curves follow a similar trend. (C) The deconvolved 3-dimentional images of the area of the acceptor cells containing MitoGFP were reconstituted with Huygens Software. These images show volumes covered by MitoGFP and TOM20 signals and their overlap.

**Supplementary Figure 3.**
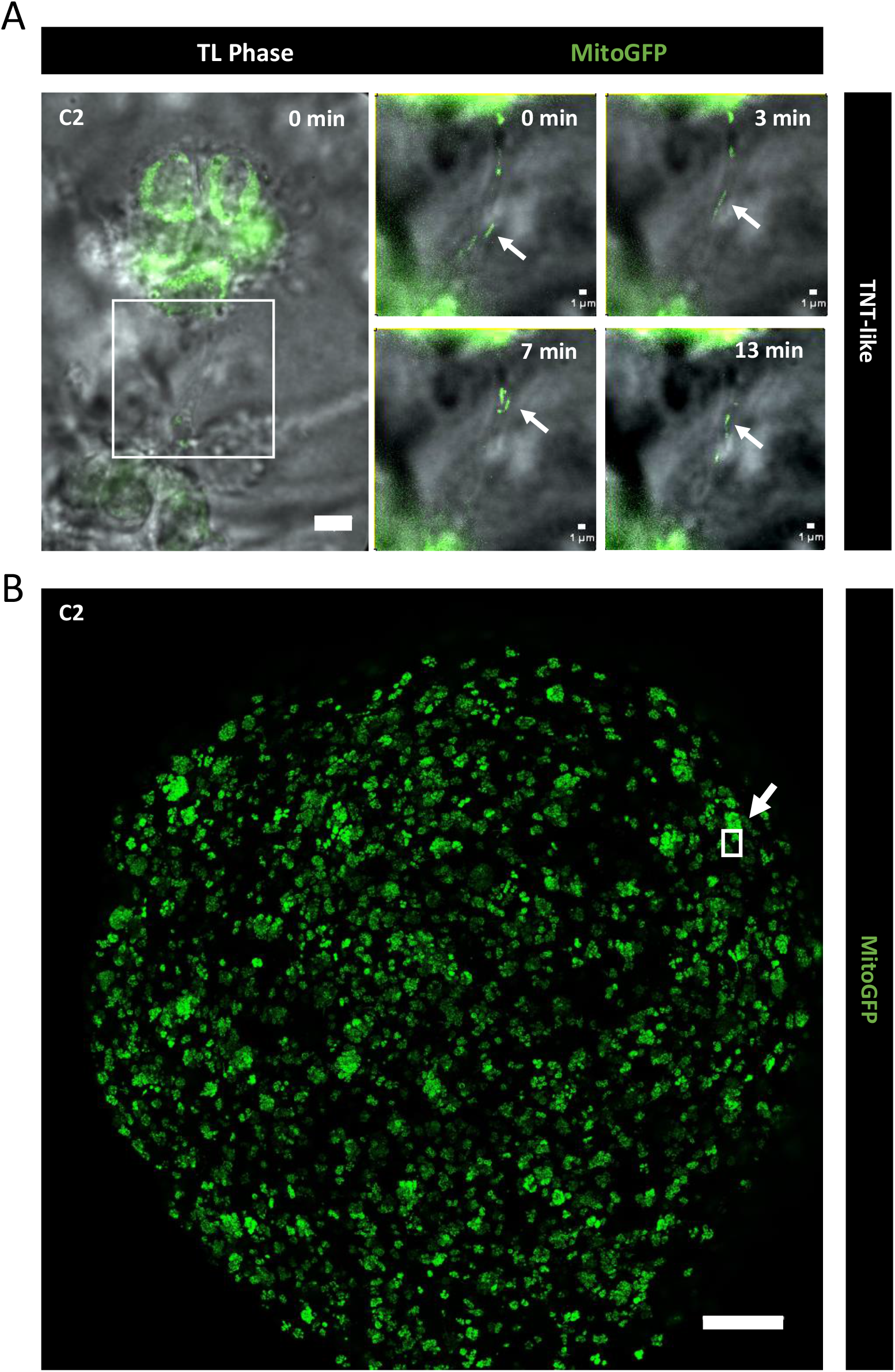
Movement of mitochondria by live-imaging in C2-tumor organoids inside TNT-like connections. (A) C2 MitoGFP tumor organoids were imaged at 7 days of culture, images composed of 25 z-stacks were acquired every 1 min for 13 min (step size 0.45 μm, total thickness ~12μm) acquired with transmitted light and green fluorescence (MitoGFP). Left panel shows an overall vision of the TNT-like connection inside the tumor organoid a time 0 min. In the right panels are shown the areas magnified at different time points from 0 up time 13 min. White arrow points at mitochondria inside the TNT-like connection. Scale bar: 10 μm. (B) 7 days-old C2 MitoGFP tumor organoids was fixed after the live-imaging and imaged by confocal microscopy. This image is the result of the z-projection of 11 stacks with 6 μm of step for a total thickness of 66 μm. The white frame with arrow is representative of the size of area where the live-imaging video was acquired, not of the specific location. Scale bar: 400 μm.

**Supplementary Figure 4.**
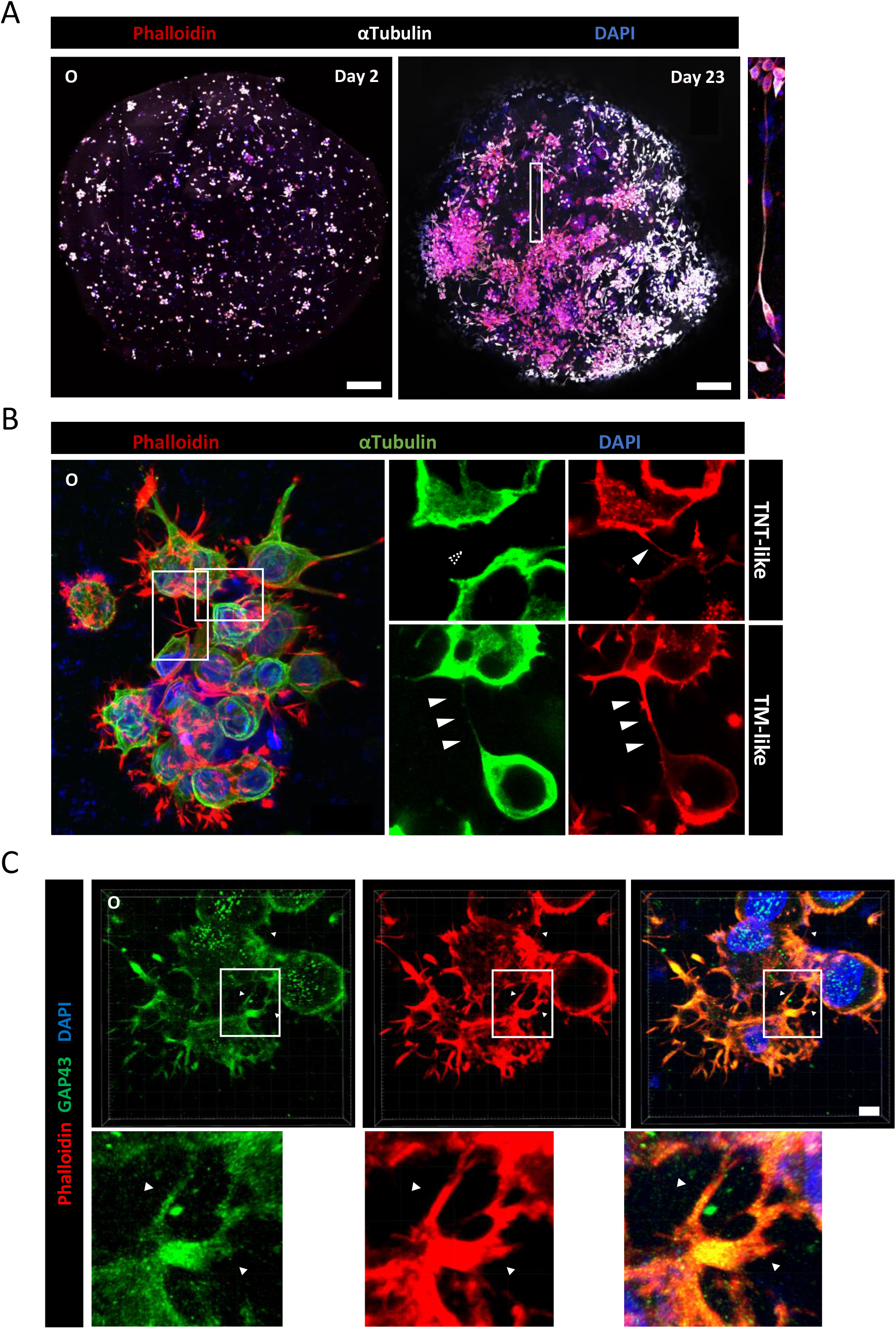
TNT- and TM-like connections in cells derived from patient O tumor. (A) Representative fluorescence images of the whole patient O-derived tumor organoid at 2 and 23 days of growth using Pln-Apo 10X/0.45 objective of inverted confocal LSM700. The resulting images represent a max intensity projection of 6 and 6 sections (step size: 5 and 4 μm), respectively, stained for anti-αTubulin (microtubules, white), Phalloidin (actin in red) and nuclei (blue). A magnification of a long TM-like extension is presented on the right panel. Scale bars are 300 (left) and 200 μm (right). (B) Representative pictures of patient O-derived organoids at 2 days, stained for anti-αTubulin (microtubules, green), Phalloidin (actin filaments, red), and nuclei (blue). Confocal images were acquired with 40X objective. Regions of interest show either αTubulin-devoid connections, defined as TNT-like (<1 μm), or thick αTubulin-positive connections (>1 μm), named TM-like. Dashed arrowheads indicate the absence of a fluorescent signal at the connection level, white-filled arrowhead show positiveness to the signal. Main image is the results of max intensity projections of 11 slices (step size: 0.5 μm), region of interest of TNT-like and TM-like connection correspond to the Z-stack 1 and 6, respectively. Scale bar: 20 μm. (C) TM-like protrusion expressing GAP43 in patient O-derived organoids. 6 days-old organoids were fixed and stained with anti-GAP43 (in green), phalloidin (actin filaments, in red) and DAPI (in blue). Confocal images with 40x objective were acquired. 3D reconstruction of a 11 sections image (step size: 0.5 μm) was performed using Imaris Viewer software. Regions of interest shows two TM-like GAP43-positive connections (>1 μm). Scale bar: 5 μm.

**Supplementary Figure 5.**
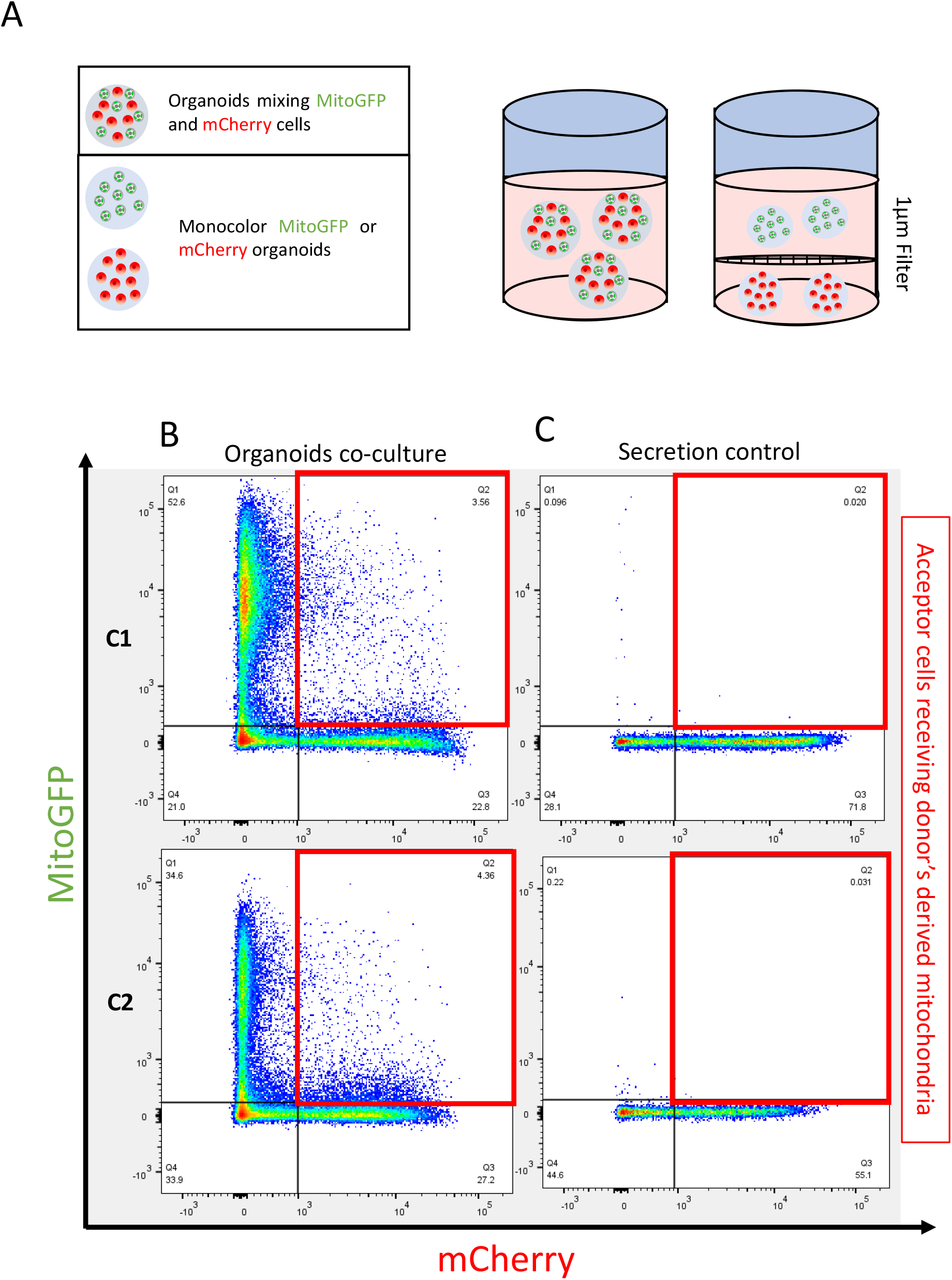
Mitochondria transfer analysis in organoids by flow cytometry. (A) Schematic of the experiment. We prepared co-culture organoids, mixing donor cells (MitoGFP) and acceptor cells (mCherry) in a 1:1 ratio, and monocolor organoids, using only donor or acceptor cells. Co-culture organoids were cultured overtime and desegregated at each timepoint to monitor the percentage of acceptor cells receiving donors’ derived mitochondria (on the left). Monocolor organoids were cultured in the same culture medium separated by a 1 μm filter (on the right). (B) Representative plot of co-culture organoids in C1 and C2 cells after 23 days of culture. GSLCs-derived co-culture organoids were prepared, individually from C1 and C2 cells, mixing in a 1:1 ratio donors cells (MitoGFP, on the Y axis) and acceptor cells (mCherry, on the X axis). After 23 days, duplicates of a pool of 3 organoids desegregated in a single cells suspension were analyzed separately by flow cytometry. Acceptor cells (mCherry) positive also for MitoGFP signal are framed in the red boxes. (C) Representative plot of secretion control in C1 and C2 cells after 23 days. Monocolor organoids, prepared of only acceptor cells (mCherry) or donor cells (MitoGFP) and cultured in the same culture medium separated by a 1 μm filter for 23 days and desegregated in a single cells suspension and analyzed by flow cytometry. Acceptor cells (mCherry) positive also for MitoGFP signal are framed in the red boxes.

**Supplementary Figure 6.**
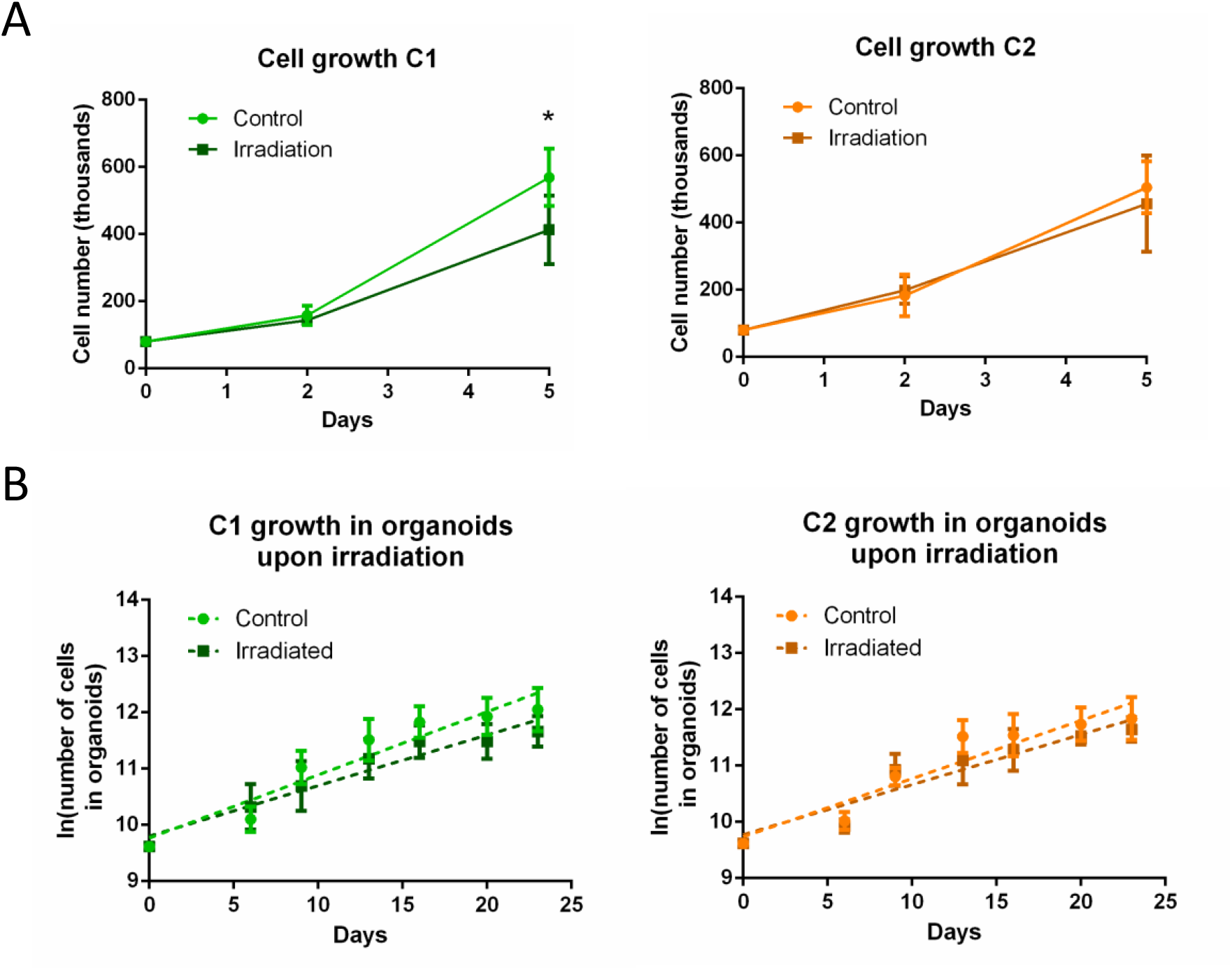
Cell proliferation after irradiation in co-culture experiments. (A) Cell growth in irradiated co-culture experiment in adherent conditions, relative to Fig. 5B. For C1, 143970± 6653 and 413000±101930 cells were counted after 2 and 5 days, respectively, in the coculture with irradiated cells (3 independent experiments). A significant reduction of cells was observed at day 5 compared to control condition (day 2: 158000±28751; day 5: 568866±85332, 3 independent experiments). For C2, 199080±40341 and 456260±143521 cells were counted after 2 and 5 days, respectively, in the co-culture with irradiated cells (5 independent experiments). No significant difference was observed compared to control at the two timepoint in analyse (day 2: 182900±61890, day 5: 505260±77515, 5 independent experiments). ANOVA two-way test was performed. P value < 0.05 (*), P values > 0.05 are not significant and not indicated on the figure. (B) Effect of irradiation on cell number in tumor organoids, relative to Fig. 5C. Irradiation was performed after 5 days from the organoid preparation. Duplicates of a pool of 3 organoids were dissociated in a single cell suspension and counted at each timepoint. Control C1: day 6 24800±5768; day 9 63150±18350; day 13 105850±43970; day 16 140450±33929; day 20 158600±60394 day 23 181800±78820. Irradiated C1: day 6 24767±I4749; day 9 47100±18499; day 13 747ÜÜ±28446; day 16 100050±32374; day 20 100700±32051; day 23 119000±29480 (4 independent experiments). Control C2:day 6 22600±3704; day 9 49700±8116; day 13 104200±33870; day 16 108580±42218; day 20 128800±34478; day 23 145080±47726. Irradiated C2: day 6 16667±6853; day 9 57150±16787; day 13 70250±29190; day 16 83400±29947; day 20 97725±10594; day 23 115600±23118 (4 independent experiments). The cell number was transformed into a logarithmic scale and slopes were compared by linear regression (dashed lines). No significant difference was observed between control and irradiated condition in both GSLCs.

**Supplementary video 1: Movement of mitochondria along TNT-like connections in 2D-conditions.** C2 MitoGFP cells were plated on laminin-coated surface and imaged after 6h. 18-z-stacks images were acquired every 1 min for 27 min (step size: 0.47 μm) with merged transmitted light and green fluorescence (MitoGFP). The movie, resulting from the max z-projection of each time-frame, shows the transfer of mitochondria between two C2 cells expressing MitoGFP and connected by TNT-like connection in 2D culture. Scale bar 10 μm.

**Supplementary video 2: Transfer of mitochondria via TNT-like connections in tumor organoids.** C2 MitoGFP tumor organoids were imaged at 6 days of culture, images composed of 62 z-stacks were acquired every 1 min for 38 min (step size 0.45 μm, total thickness ~28 μm). On the left panel, video corresponding to the merge of time-frame images acquired with transmitted light and green fluorescence (MitoGFP). Only green fluorescence image is shown in the right panel, for better visualization. Videos are resulting from the max-z-projection. White and red arrows point at the movement of mitochondria inside TNT- and TM-like connections, respectively. Scale bar 10 μm.

**Supplementary video 3: Motion of mitochondria inside TNT-like connections in tumor organoids.** C2 MitoGFP tumor organoids were imaged at 7 days of culture, images composed of 25 z-stacks were acquired every 1 min for 13 min (step size 0.45 μm, total thickness ~12μm). Video corresponds to the merge of time-frame images acquired with transmitted light and green fluorescence (MitoGFP). White arrow points at mitochondrion inside the TNT-like connection. Scale bar 10 μm.

## Notes

### Competing Interest Statement

The authors have declared no competing interest.

